# Immunomodulatory impact of *PTPN11*/SHP2-based vertical RAS-MAPK pathway inhibition in pancreatic cancer

**DOI:** 10.64898/2026.05.23.727401

**Authors:** Asma Y. Alrawashdeh, Xun Chen, Philipp Hafner, Steffen J. Keller, Tonmoy Das, Huda Jumaa, Thomas Y. Avery, Solène Besson, Silke Hempel, Stephanie Mewes, Kerstin Meyer, Mohammad Al Shhab, Mara Schneider, Johannes D. Lettner, Dominik von Elverfeldt, Wilfried Reichardt, Melanie Boerries, Stefan Fichtner-Feigl, Geoffroy Andrieux, Dietrich A. Ruess

## Abstract

Protein tyrosine phosphatase non-receptor 11 (*PTPN11*/SHP2) is a critical upstream mediator of RAS-MAPK signaling and a central node in adaptive resistance mechanisms evolving with RAS and MEK/ERK inhibition. Accordingly, clinical trials are currently evaluating allosteric SHP2 inhibitors in vertical RAS pathway combination therapies for various KRAS-mutant malignancies, including pancreatic ductal adenocarcinoma (PDAC).

Here, we aimed to delineate the immunomodulatory effects of SHP2-based vertical RAS pathway inhibition in notoriously immunotherapy-refractory PDAC, spanning from early treatment response to invariably evolving adaptive resistance. Employing human and murine *PTPN11* knockout and wild-type PDAC cell lines, an autochthonous murine PDAC model (KPC), and patient-derived PDAC organoids, we find that short term dual MEK/SHP2 inhibition induces increased T cell infiltration and a reduction in immunosuppressive M2-like macrophages. However, these effects are accompanied by a decrease in mature dendritic cells and a concomitant expansion of monocytic myeloid-derived suppressor cells, indicative of a mixed immunological response with both immune-activating and immune-suppressing features. These changes are associated with tumor cell-intrinsic upregulation of CXCR3 ligands and TGF-β, as well as increased expression of checkpoint ligands for TIGIT and TIM-3 across molecular subtypes and species. With prolonged treatment and transition to an adaptive resistant tumor cell state, the initial immune-sensitizing effects are lost and the immune-suppressive features prevail. M2-like macrophages re-accumulate, dendritic cell maturation remains impaired, TGF-β expression persists, and TIGIT and TIM-3 ligand expression is further enhanced. Notably, dual SHP2/RAS inhibition recapitulates the observed induction of TGF-β and checkpoint ligands.

Collectively, these findings identify a dynamic but ultimately immunosuppressive remodeling of the tumor microenvironment in response to SHP2-based vertical RAS pathway inhibition in PDAC and provide a rationale for combinatorial immunotherapy strategies. In particular, concurrent targeting of TGF-β, combined TIGIT/TIM-3 checkpoint blockade, and likely CD40 agonism may help sustain early immune activation while counteracting emerging suppressive features, thereby improving the durability of tumor control.

## Introduction

PDAC accounts for over 90% of pancreatic malignancies^1^ and is projected to become the second leading cause of solid tumor related mortality by 2040 in many high-income countries^2^. While surgical resection in conjunction with (neo-) adjuvant chemotherapy remains the most effective and the only potentially curative strategy, only 10–20% of patients are diagnosed at an early stage, with resectable disease^3^. Therefore, systemic (chemo-) therapy remains a cornerstone for PDAC treatment in both (neo-) adjuvant and palliative settings^4^. The introduction of multiagent chemotherapeutic regimens has led to some improvement of chemotherapeutic efficacy and patient prognosis, e.g. FOLFIRINOX (5-fluorouracil, leucovorin, irinotecan, and oxaliplatin) and the combination of gemcitabine with nanoparticle albumin-bound paclitaxel (nab-paclitaxel) both showing improved overall survival of patients with metastasized disease in comparison to the historical standard-of-care gemcitabine monotherapy^5,6^. Also, as a consequence, about 20% of patients with primarily unresectable non-metastatic PDAC may today undergo surgical exploration and resection, i.e. conversion surgery, following 4-6 months of induction combination chemotherapy with or without radiotherapy^7^. But even with these recent developments PDAC remains associated with a dismal prognosis, with a 5-year overall survival rate of approximately only maximally 13%^8^, and there is great need for additional innovative therapeutic options.

Due to its strongly desmoplastic stroma and immunosuppressive tumor microenvironment (TME), its low mutational burden and low frequency of deficient DNA mismatch repair resulting in a low number of neoepitopes, immunotherapeutic approaches in PDAC have been rather disappointing. Oncogenic KRAS mutations, which are observed in approximately 95% of PDAC cases^9^ and are a key driver of proliferation, metabolic adaptation, survival and immune evasion, have always been considered a promising target for therapeutic intervention. Until recently, only substances targeting downstream effectors of KRAS, such as MEK1/2 or ERK1/2, had been available, but largely failed to demonstrate improved outcomes, often due to receptor tyrosine kinase (RTK)-dependent feedback-activation, resulting in reestablishment of downstream signaling^10,11^. A plethora of direct RAS and KRAS inhibitors is now revolutionizing the field, with many compounds being at the threshold for or having already reached clinical application^12^. Yet, resistance emerges rapidly with these drugs as well, and combination approaches will be necessary for sustained efficacy and patient benefit.

Previously, we and others found that SHP2 is a critical and positive upstream mediator of mutant KRAS signaling output and a central node in adaptive resistance mechanisms evolving with MEK/ERK inhibition^13–16^. Accordingly, clinical trials are underway evaluating allosteric SHP2 inhibitors in vertical RAS pathway combination therapies for various KRAS-mutant malignancies, including PDAC^17^. It has been reported that SHP2 inhibition, alone or in combination with KRAS^G12C^ inhibition, may remodel the TME of KRAS^G12C^-driven non-small cell lung cancer (NSCLC) or PDAC in an orthotopic transplantation model, decreasing myeloid suppressor cells, increasing CD8+ T cells, and sensitizing to PD-1 blockade^18^. Importantly, beyond tumor cells, the SHP2/RAS/ERK pathway is also critical for immune cell homeostasis^19^. T cell priming, expansion and memory require accurate modulation of signaling along this axis, and the role of SHP2 extends to involvement in propagation of programmed cell death 1 (PD-1) and B- and T-lymphocyte attenuator (BTLA) immune checkpoint signals^20^, not restricted to T- and B-cells but also e.g. including mechanisms of PD-1-SHP2 mediated myelocyte differentiation^21^.

Here, we sought to understand the immunomodulatory consequences of continuous SHP2-targeted vertical RAS pathway inhibition in immunotherapy-refractory PDAC, covering both early treatment response and the development of invariably evolving adaptive resistance. We further aimed to establish a molecular rationale to inform the selection of future combinatorial immunotherapy agents to improve the durability of tumor control.

## Results

### Vertical RAS pathway inhibition with a combination of SHP099 and trametinib delays PDAC progression *in vivo*

The autochthonous *Kras*^LSL-G12D/+^; *Trp53*^fl/fl^; *Ptf1a*^Cre-ex1/+^ (KPC) PDAC model spontaneously develops pancreatic tumors which recapitulate human disease heterogeneity and tissue context, enabling the study of tumor progression, therapeutic response and immune microenvironment alterations in a translationally meaningful way. KPC mice were monitored with MRI for tumor occurrence starting at approximately four weeks of age and, upon PDAC detection, were subjected to a short-term treatment regimen and analysis (2 weeks), or long-term treatment and overall survival (long-term) analysis; the full experimental timeline is outlined in (**Fig. 1a**). Mice randomly received either SHP099, trametinib, a combination of both, or a vehicle control. Tumor volume dynamics were assessed weekly by MRI.

**Fig. 1.**
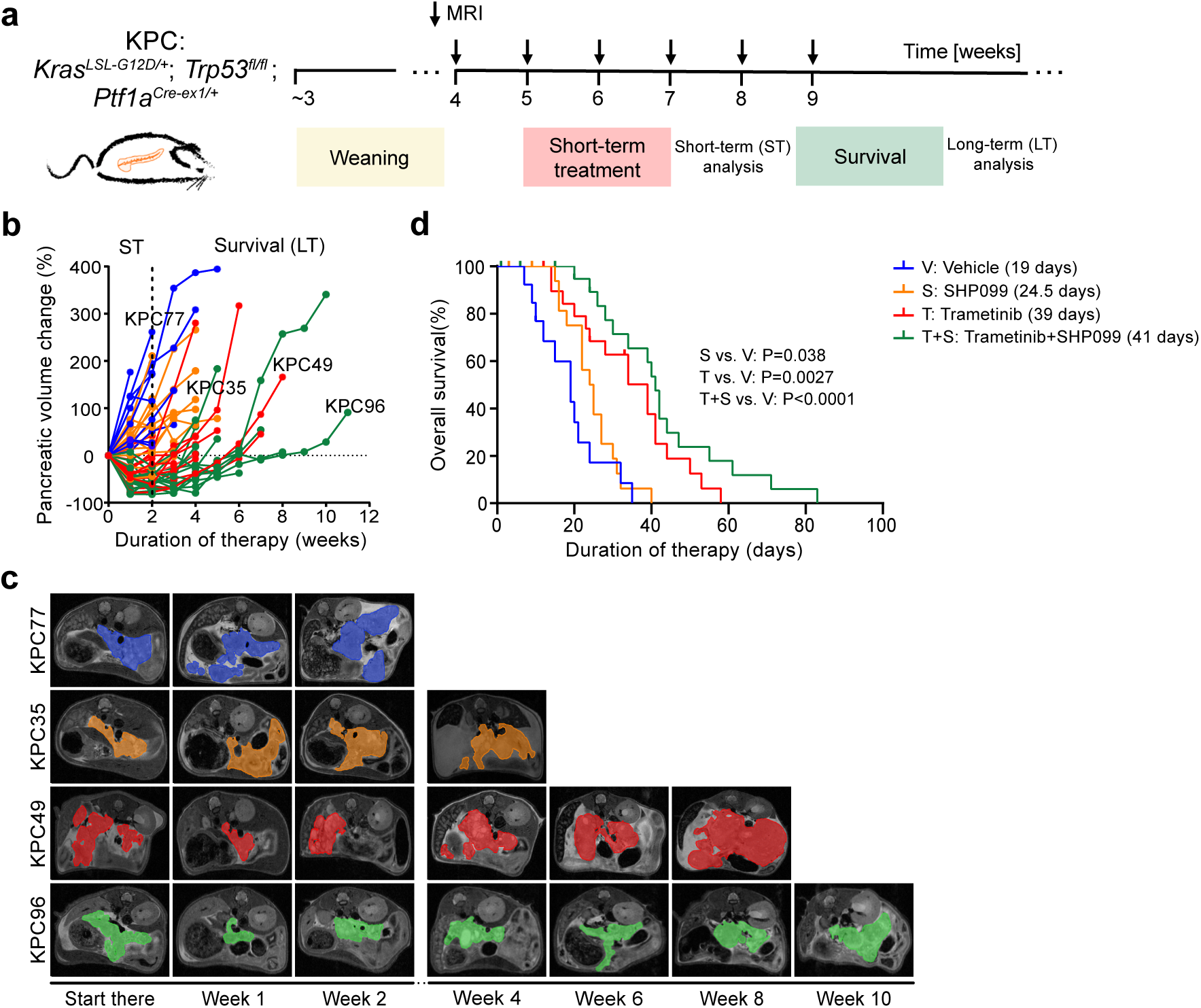
Vertical RAS pathway inhibition with a combination of SHP099 and trametinib delays PDAC progression *in vivo*. **a.** Experimental timeline for in vivo treatment and analysis in KPC mice. **b.** Quantification of pancreatic volume change over time following detection as measured by MRI. V: Vehicle cohort (n= 9), S: SHP099 cohort (n=16), T: trametinib cohort (n=14), or T+S: trametinib+SHP099 cohort (n=14). The dotted line marks the baseline tumor volume at the short-term (ST) treatment time point (2 weeks). **c.** Representative MRI scan slices depicting PDAC tumor sections of four KPC mice treated with V (KPC77), S (KPC35), T (KPC49) or T+S (KPC96) at the indicated time points (weeks) following the start of therapy (start ther). **d.** Kaplan–Meier analysis of tumor-related survival in KPC mice treated with V (n=13), S (n=17), T (n=19) and T+S (n=21). Ticks represent censored mice euthanized due to a decline in clinical condition in the absence of histological evidence of frank PDAC. Significance was determined by log-rank (Mantel–Cox) test. Median survival is indicated in brackets.

Unlike GS493, a catalytic site SHP2 inhibitor with pharmacokinetic and -dynamic shortcomings which had shown no significant monotherapy benefit compared to vehicle in own prior work^13^, the allosteric SHP2 inhibitor SHP099 significantly prolonged survival in comparison to vehicle controls (24.5 vs. 19 days; *P* = 0.038) as monotherapy, although it did not lead to tumor regressions (**Fig. 1b-d**). Trametinib was confirmed to potently achieve an initial reduction in pancreatic tumor volume; however, by about week 4, tumor volumes rebounded to nearly 100%, and continued to progress afterward, indicating acquired resistance. In contrast, the addition of SHP099 to trametinib delayed resistance development, maintaining pancreatic tumor volumes below baseline (0% volume change) up to week 6, reflecting sustained suppression of tumor progression (**Fig. 1b**). Representative MRI scans from four individual mice illustrate tumor burden across treatment groups, with dual therapy-treated mice exhibiting notably slower tumor progression and smaller tumor volumes between weeks 4-8 compared to monotherapies or vehicle (**Fig. 1c**). Accordingly, KPC mice receiving trametinib alone had a median survival of 39 days, while those in the combination group had a median survival of 41 days and survived up to ∼80 days (**Fig. 1d**).

These data demonstrate that the addition of allosteric SHP2 inhibition to MEK inhibition, within a fully oral targeted combination regimen, enhances therapeutic efficacy in PDAC by delaying adaptive resistance and improving overall survival in an aggressive endogenous tumor model.

### Dual MEK/SHP2 inhibition remodels the PDAC immune microenvironment, transiently enriching pro-inflammatory immune subsets

We then sought to characterize immunomodulatory effects of SHP2-based vertical inhibition of the RAS pathway using the aforementioned KPC mouse trial, encompassing early treatment response through the emergence of adaptive resistance. Immune cell subsets were profiled by flow cytometry (FC) and corroborated using multiplex immunohistochemistry (mIHC).

FC analysis of CD45⁺ immune cells isolated from PDAC tumors following short-term (2-week) treatment revealed a tendency of increased CD4⁺ T cell numbers in the dual therapy group but there were no notable changes in Th1 or regulatory T cell (Treg) populations. CD8^+^ cytotoxic T cells, including activated subsets (CD25^+^), as well as γδ T cells and NK cells, remained largely unchanged. We observed a potentially beneficial alteration with respect to tumor-associated macrophages, marked by a shift toward a pro-inflammatory state, with increased F4/80⁺CD86⁺ (M1-like) macrophages and a significant decrease (p < 0.005) in F4/80⁺Arg1⁺ (M2-like) macrophages, resulting in a higher M1/M2 ratio (p < 0.05). In parallel, monocytic myeloid-derived suppressor cells (mMDSCs) were significantly elevated in the dual therapy group, whereas granulocytic MDSCs (gMDSCs) showed a decreasing trend. The proportion of mature dendritic cells (mDCs) declined, while conventional dendritic cell type 2 (cDC2) populations remained unaffected. However, and importantly, all significant effects observed with short-term treatment were lost in the long-term treatment cohorts, coinciding with disease progression and the emergence of adaptive treatment resistance (**Fig. 2, Fig. S1a)**. Gating strategies are disclosed in (**Fig. S1b)**.

**Fig. 2.**
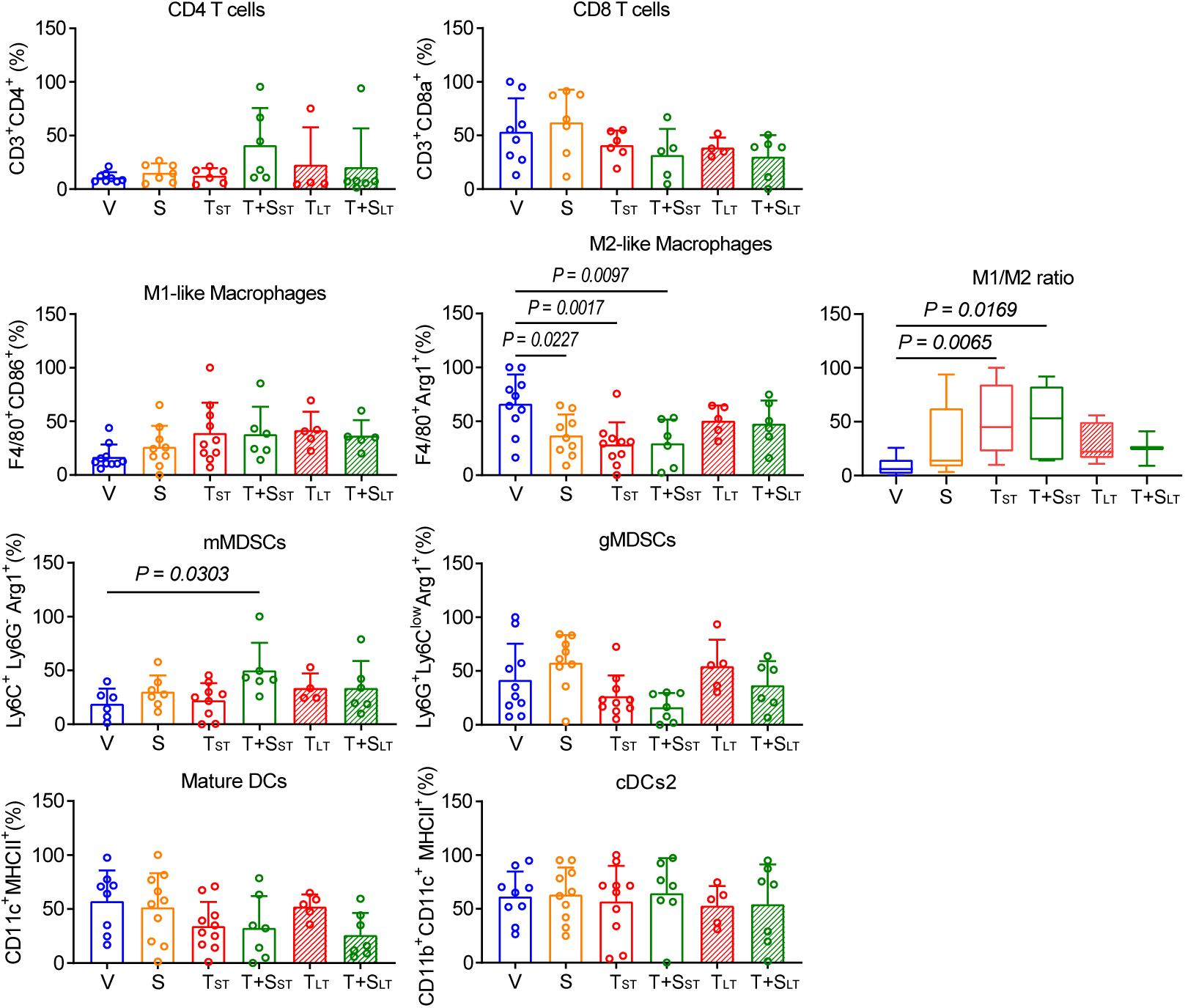
Dual MEK and SHP2 inhibition diminishes immunosuppressive macrophages in PDAC. Flow cytometry–based quantification of immune populations across PDAC samples, including T cells, macrophages, myeloid-derived suppressor cells (MDSCs), and dendritic cells (DCs). Data are shown for each treatment group at short-term (2 weeks) and survival endpoints. A reduction in anti-inflammatory (M2-like) macrophages is reflected by changes in the M1/M2 ratio. Statistical analysis was performed using one-way ANOVA across the cohort; bars represent mean ± SD. Significance is indicated as *p* < 0.05 and *p* < 0.005. Frequencies are shown relative to the CD45⁺ population.

To validate and contextualize these FC findings, we employed mIHC on OPAL-stained FFPE tissue sections from 74 murine samples for this study. Three whole tumor tissue sections were stained for each single mouse employing three different mIHC panels: a Lymphoid cell panel (CD8^+^ T cells, CD4^+^ T cells, NK T cells, and Tregs), a Myeloid cell panel (macrophages, monocytic- and granulocytic-MDSCs), and a Dendritic cell panel (mature, resident and migratory dendritic cells), (exemplified by **Fig. 3a**, top panel). We quantified tumor immune cells across whole-organ tissue sections. Over time, with both long-term trametinib and trametinib+SHP099, the overall density of lymphoid and dendritic cells appeared to decrease, whereas myeloid cells, primarily represented by macrophages, remained dominant across treatment arms and durations (**Fig. 3a**).

**Fig. 3.**
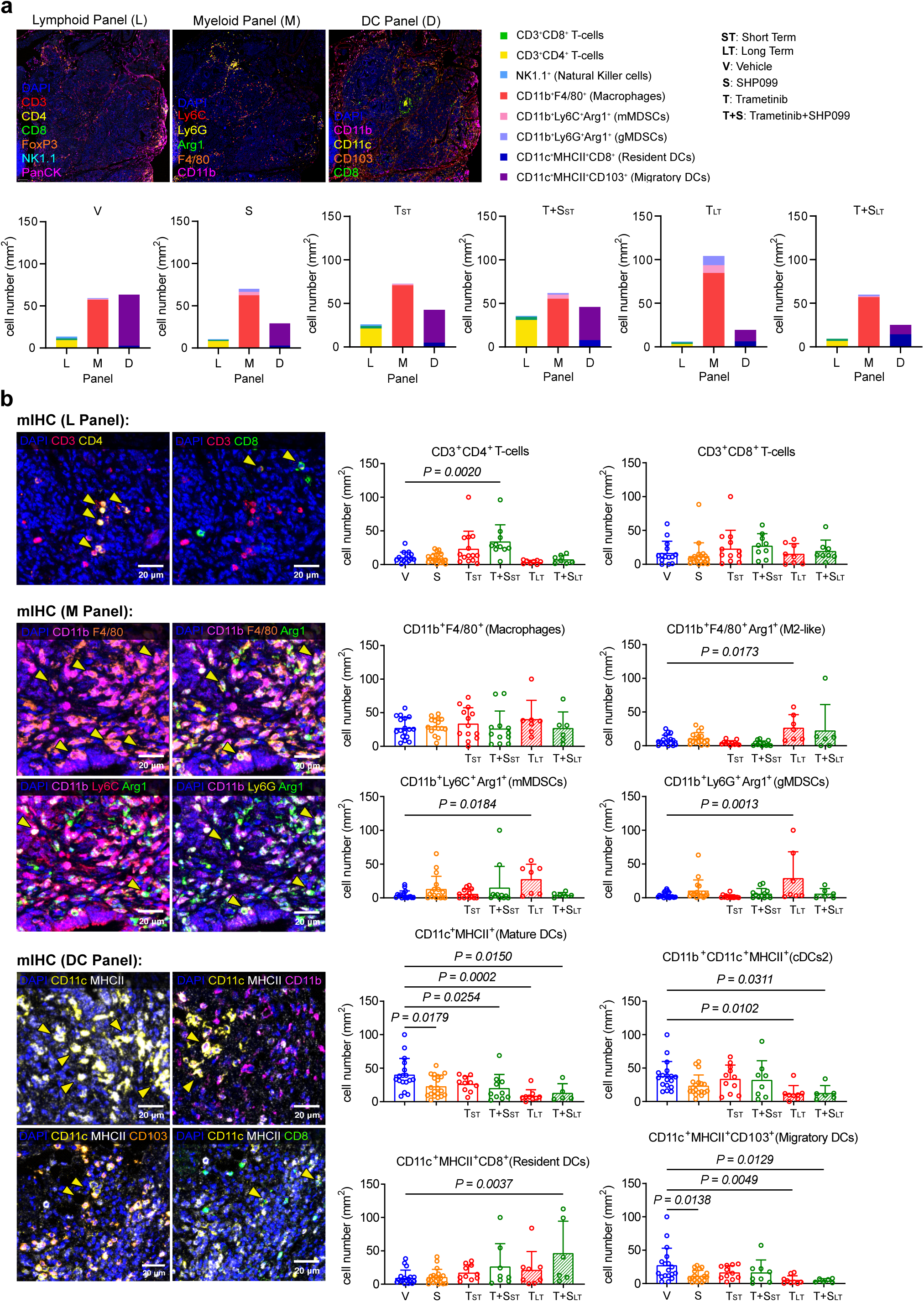
Multiplex IHC reveals the broader immune context of dual MEK and SHP2 inhibition in PDAC. FFPE sections of PDAC tumors from the four treatment arms **a.** were stained using OPAL multiplex immunofluorescence for three immune panels: the “Lymphoid cell panel,” including PanCK (epithelial marker), CD3, CD4, CD8, NK1.1, and FOXP3; the “Myeloid cell panel,” including CD11b, F4/80, arginase, Ly6G, and Ly6C; and the “Dendritic cell panel,” including CD11c, MHCII, CD103, CD8, and CD11b. Whole-tissue sections were imaged at 20× magnification using the Akoya PhenoImager HT system. Immune context quantification from the three panels was performed and plotted in GraphPad Prism as cell number per mm² for each treatment arm at both short-term and survival time points. **b.** Collated data for all PDAC samples in the study cohort are shown, including cell number per mm² of T cells (CD3⁺), myeloid cells (CD11b⁺), and dendritic cells (CD11c⁺). Statistical analysis was performed across the cohort using one-way ANOVA; bars represent mean ± SD, and significant differences are indicated as p < 0.05, p < 0.005, and p < 0.0005.

In more detail, we observed three key changes (**Fig. 3b**): (1) a significant increase in CD3⁺ T cells, particularly the CD3⁺CD4⁺ subset, with dual therapy in short-term treatment, aligning with FC data; (2) elevated populations of CD11b⁺ myeloid cells in the long-term trametinib arm, which were reduced by SHP099 addition (i.e. long-term dual inhibition), including M2-like macrophages, monocytic MDSCs (mMDSCs), and granulocytic MDSCs (gMDSCs); and (3) a marked reduction in dendritic cell subsets in long-term treatment arms, with mature (MHCII^+^) DCs being strongly affected, which included CD103^+^ cDC1-like and Cd11b^+^ cDC2s. Notably, CD8a^+^ resident DCs significantly increased in KPC tumors with long-term dual MEK/SHP2 inhibition, yet with high variability across individual samples. Treg and NKT cells showed no notable changes in mIHC analysis, consistent with our FC findings (**Fig. S2a**). Single-channel images are shown in (**Fig. S2b**).

We aimed to further validate by examining long-term treated PDAC tissue sections from our historical GS493 and trametinib cohort^13^. Here, dual inhibition including GS493 resulted in a sustained long-term decrease in the M2-like macrophage phenotype, unlike with SHP099. For DCs, however, GS493 showed a decreasing trend consistent with observations from SHP099-treated samples. Since this historical cohort of GS493-treated mice were handled at a different institution, not only distinct pharmacokinetics and -dynamics, but also factors such animal housing and microbiota composition could have contributed to the observed differences (**Fig. S3a,b**).

While mIHC generally provides more reliable quantification of myeloid and dendritic cell populations, which are susceptible to loss or phenotypic alteration during tissue dissociation required for FC, the two approaches together provided complementary insights into the immune landscape following dual trametinib and SHP099 treatment in KPC PDAC. Our data indicate that short-term MEK or MEK/SHP2 inhibition induces a mixed immune response characterized by both immunostimulatory and immunosuppressive features. In contrast, prolonged treatment promotes re-establishment of an immunosuppressive tumor microenvironment, which is only partially attenuated by including SHP099. This effect was particularly evident within the myeloid compartment, where M2-like macrophages showed marked reaccumulation in both long-term treated cohorts.

### Proteomic screening for immune-related changes in murine KPF and human PDAC cell lines *in vitro*

To screen for cancer cell-intrinsic alterations induced by pharmacologic and genetic interference with SHP2, with or without MEK inhibition, that may regulate immune cell crosstalk and trafficking, we employed previously established dual recombinase KPF murine tumor cell lines that allow a Tamoxifen-inducible ex vivo SHP2/PTPN11 genetic deletion (F2453 and F2683; *Kras^FSF-G12D/+^; Trp53^frt/frt^; Ptf1a^Flpo/+^; R26^FSF-CreERT/+^; Ptpn11^fl/fl^*)^13^ (**Fig. S4a,b**). First, we evaluated and confirmed on target effects via signaling readout by western blot. As expected, genetic or pharmacologic inactivation of SHP2 meaningfully suppressed phosphorylated ERK (pERK) in the context of MEK inhibition, which alone, was associated with increased KRAS^G12D^ expression, reactivation of MEK (increased pMEK levels) and parallel signaling pathway activation (pAKT). pSTAT3 responses were cell line–dependent (**Fig. S4a,b**). We then assessed the impact of dual inhibition on cytokine and chemokine levels in tumor cell lysates, employing commercial cytokine arrays (**Fig. S4c-h**). In F2683, dual treatment induced ICAM-1, cystatin C, and IL-1Ra. In F2453, dual inhibition broadly reduced cytokines, including osteopontin (OPN), VCAM-1, CX3CL1, and CXCL5, which may be linked to pro-tumorigenic signaling and immune suppression in pancreatic cancer^22–24^ (**Fig. S4c,d**). However, overall, no truly significant differences in total cytokine or interleukin levels were observed in this exploratory experiment (**Fig. S4e,f**).

In a parallel, second screen, we performed mass spectrometry-based proteomic analyses on 3 human cell lines (YAPC, PANC-1, and MIA PaCa-II; WT vs. *PTPN11* deleted by CRISPR/Cas9) treated for 72 hours (short-term). First of all, SHP099-mediated pharmacologic inhibition revealed an only minor and strongly discrepant impact on the proteome, compared to *PTPN11* genetic deletion, precluding meaningful deductions from the knockouts. Single agent trametinib treatment strongly altered the proteome, which was further pronounced with the addition of SHP099 in the context of dual inhibition (**Fig. S5a**). With dual inhibition, proteins associated with innate immune response and response to external stimulus were positively enriched in MIA PaCa-II and YAPC cells, whereas PANC-1 cells showed no significant changes (q < 0.05; data not shown for PANC-1). Notably, Type I interferon-mediated signaling and cytokine-mediated signaling pathways were specifically enriched in YAPC cells (**Fig. S5b**).

### Transcriptional profiling reveals distinct biological and immune-driven programs associated with treatment response and molecular subtypes in PDAC

With the aim to further clarify the molecular underpinnings of tumor-immune cell crosstalk and trafficking we then performed RNA sequencing across a spectrum of preclinical models to investigate subtype-specific differences between short- and long-term responses to vertical RAS-MAPK pathway inhibition, including the KRAS^G12D^ mutant specific inihibitor MRTX1133, on a broader scale. Specifically, we analyzed transcriptomic profiles from 14 KPC cell lines and 3 human pancreatic cancer cell lines, representative of the molecular subtype spectrum of PDAC. To model treatment resistance, human and selected KPC cell lines were continuously exposed to fixed drug concentrations until they resumed proliferation. All samples were compared to their respective DMSO-treated controls. We assessed fast gene set enrichment (fgsea) in Hallmark (H) gene sets, Gene Ontology (GO) biological processes, Kyoto Encyclopedia of Genes and Genomes (KEGG) pathways, and BioCarta pathways. Transcriptomic analysis identified five major functional categories: (1) Cell Cycle, (2) Tissue Remodeling, (3) Oncogenic Signaling, (4) Immune Response, and (5) Migration/Chemotaxis (**Fig. 4a-c**). To complement our pathway-level insights obtained through fgsea, we performed a comprehensive differential gene expression (DGE) analysis using a curated set of immune-related genes from literature and our dataset.

**Fig. 4.**
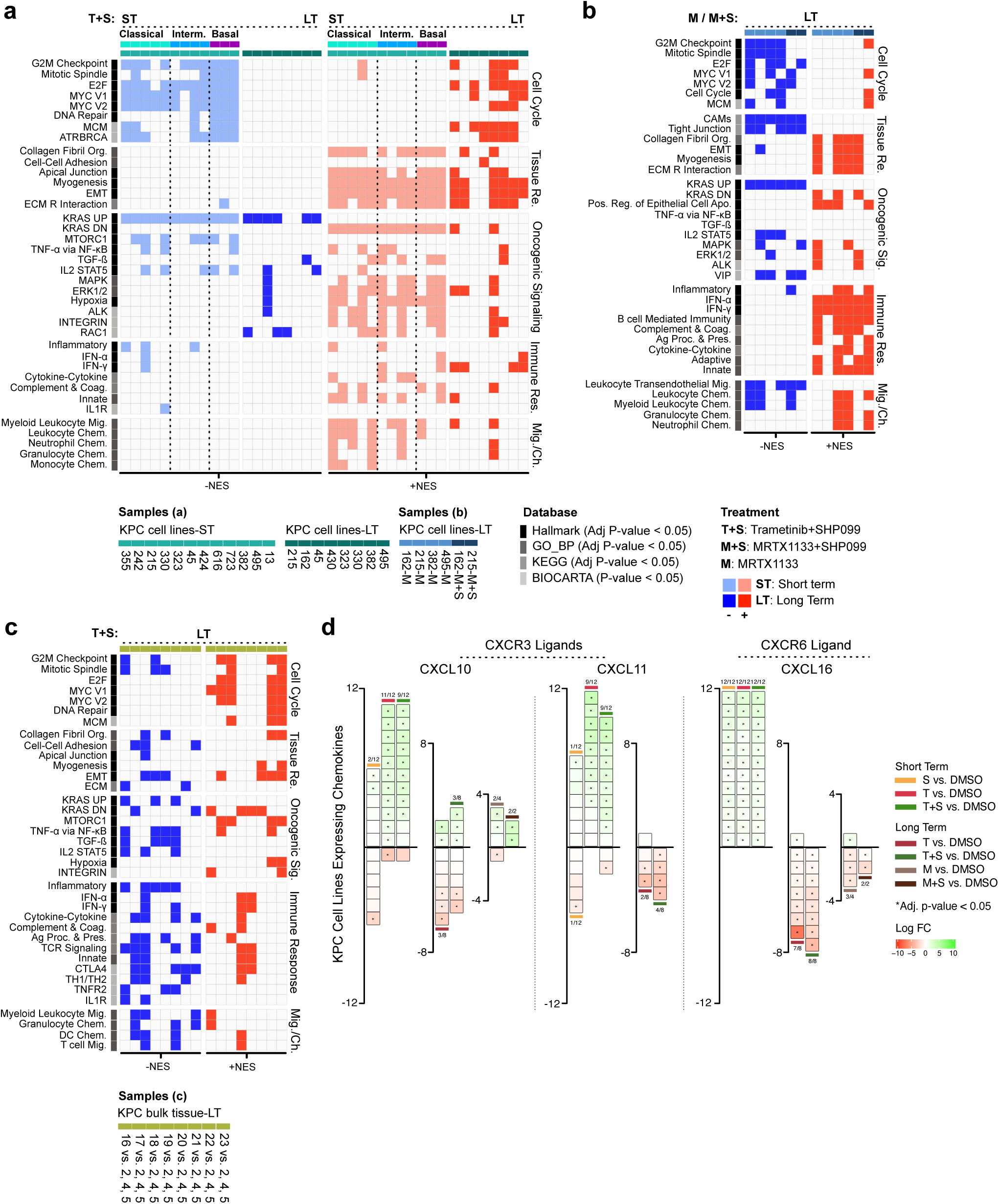
Transcriptional profiling reveals distinct biological and immune-driven programs associated with treatment response and molecular subtypes in PDAC. Gene set enrichment analysis (GSEA) of significantly enriched Hallmark, GO_BP, KEGG (adjusted p < 0.05), and BIOCARTA (p < 0.05) pathways, grouped into five functional categories: cell cycle regulation, tissue remodeling, oncogenic signaling, immune response, and migration/chemotaxis. Normalized enrichment scores (NES) are shown (blue, downregulated; red, upregulated). Sample type, database, treatment, and duration are indicated below. Treatment durations: ST, short term; LT, long term. Treatments: S, SHP099; T, trametinib; T+S, trametinib + SHP099; M, MRTX1133; M+S, MRTX1133 + SHP099. **a.** KPC-derived cell lines treated with T+S under ST and LT conditions; PDAC subtypes (Classical, Intermediate, Basal) indicated above. **b.** KPC-derived cell lines treated with M or M+S under LT conditions. **c.** KPC bulk tumor tissue treated with T+S under LT conditions. **d.** DEGs across KPC cell lines. Color intensity indicates log fold change (logFC; red, downregulated; green, upregulated). Asterisks (*) denote significant differential expression (adjusted p < 0.05). Numbers indicate the fraction of samples with significant changes per treatment arm.

KPC-derived cell lines were established from individual PDAC tumors (one per KPC mouse) and classified into classical, intermediate, and basal molecular subtypes based on ssGSEA-derived enrichment scores from published PDAC subtype gene sets^25–29^. Subtype assignment was further validated by western blot analysis of representative marker proteins (**Fig. S6a,b**). E.g. the treatment-naïve parental cell lines 382, 495, and 13 exhibited reduced levels of epithelial markers, including E-cadherin and GATA6, alongside increased expression of mesenchymal markers like N-cadherin, Slug, and Snail, supporting their classification as basal-like. In parallel, KRAS^G12D^ protein levels and GTP loading were elevated predominantly in these cell lines, further reinforcing this classification. In contrast, lines with higher epithelial and lower mesenchymal marker expression were classified as classical (**Fig. S6b**). This panel enabled analysis of subtype-stratified transcriptional responses across genetically distinct backgrounds under short- and long-term conditions.

As expected, cell cycle (category 1) and oncogenic signaling (category 3) together defined a low-proliferative state in short-term dual-treated lines, most prominently in basal-like subtypes. This was evidenced by downregulation of KRAS_UP and lack of E2F and MYC target activation (Adj. p <0.05), key drivers of proliferation and oncogenesis^30^. In contrast, long-term–treated lines demonstrated reactivation of key regulatory pathways (e.g., MCM, ATRBRCA), which restore cell cycle progression and DNA repair capacity^31^, respectively (**Fig. 4a**). Notably, long-term treatment of KPC cell lines with MRTX1133, either alone or with SHP099, better maintained suppression of cell cycle programs (**Fig. 4b**).

Category 2 (tissue remodeling) largely persisted in dual-treated cells, except collagen fibril organization, which was strongly upregulated short term but decreased over time across subtypes. EMT remained upregulated in most short- and long-term–treated cells regardless of subtype (**Fig. 4a**). Similarly, MRTX1133-treated cells, alone or in combination, showed consistent upregulation of EMT signature pathways over time (**Fig. 4b**). Within category 3 (oncogenic signaling), short-term dual-treated cell lines showed heterogeneous downregulation of mTORC1 and IL2/STAT5 signaling independent of molecular subtype, while TGF-β was preferentially upregulated in intermediate and basal-like lines. Together, these patterns indicate a shift toward an adaptive, immunosuppressive, EMT-associated state (**Fig. 4a**).

Category 4 showed a dominant innate immune signature in 50% (6/12) of short-term dual-treated lines, mainly in the classical subtype. Category 5 (migration/chemotaxis) showed strong upregulation of myeloid leukocyte migration and chemotaxis pathways, also enriched in classical lines, with more than half of the cell lines (7/12) reaching significance (Adj. p <0.05). This enrichment was reduced under long-term treatment (**Fig. 4a**). Overall, trametinib monotherapy induced changes similar to dual therapy, but more strongly activated cytokine and innate immune signatures, along with increased migration/chemotaxis particularly in classical and intermediate short-term settings. SHP099 showed comparable pathway trends but was insufficient to suppress proliferation and instead promoted TNFα/NF-κB (**Fig. S6c,d**). In MRTX1133-treated lines, either alone or with SHP099, immune response pathways were robustly upregulated, including all interferon-α (IFN-α) and interferon-γ (IFN-γ) pathways (6/6) and a subset of inflammatory and cytokine interaction pathways (3/6) (**Fig. 4b**). Collectively, these findings indicate that KRAS/SHP2 inhibition recapitulates and in part enhances MEK/SHP2-driven transcriptional shifts, with MRTX1133 consistently amplifying interferon signaling and chemokine-mediated immune cell recruitment.

Among immune-related DEGs, chemokines were most consistently regulated, particularly those involved in T cell recruitment, prompting focused analysis of chemokine-mediated immune cell trafficking. Dual MEK/SHP2 inhibition induced an immune sensitization signature characterized by CXCR3 ligands (Cxcl10, Cxcl11) and the CXCR6 ligand Cxcl16^4^. In short-term dual-treated cells, Cxcl10 and Cxcl11 were upregulated in 75% (9/12) of cases, while Cxcl16 was universally upregulated (12/12). In contrast, this response was largely lost with long-term treatment, with Cxcl10 only retained in 3/8 (trametinib+SHP099) and 2/2 (MRTX1133+SHP099) conditions, and complete loss of Cxcl11 and Cxcl16 (0/8 each) (**Fig. 4d**). Collectively, these results indicate a transient interferon-linked chemokine program driving CXCR3/CXCR6-mediated T cell recruitment in the context of short-term treatment.

We then extended fgsea to human PDAC cell lines representing distinct molecular subtypes, including YAPC (classical), PANC-1 (intermediate), and MIA PaCa-II (basal-like) (**Fig. S5c**), in all of which short-term dual treatment reduced proliferation (**Fig. S5d**). Only YAPC showed upregulation of categories 2–5 in the short-term setting, consistent with previously observed proteomic induction of interferon, cytokine, and innate immune programs, in line with its classical subtype (**Fig. S5b,d**). In contrast, long-term treatment induced broad transcriptional upregulation of immune response and migration/chemotaxis in all cell lines (**Fig. S5d**). Accordingly, all models showed increased CXCR3/CXCR6 ligands, mainly CXCL10 and CXCL11, in resistant states (**Fig. S7a**).

Although human PDAC cell lines in contrast showed more sustained induction of CXCL10/CXCL11, particularly in YAPC and resistant states, these data together suggest CXCR3/CXCR6 ligand–mediated T cell recruitment contributing to the observed immune-related phenotype in KPC mice.

### Adaptive resistance to prolonged MEK/SHP2 inhibition in KPC tumors *in vivo* is accompanied by immune evasion signatures

To connect our *in vitro* proteomic and transcriptomic screening with the *in vivo* tumor context, we performed bulk RNA sequencing of tumor tissues from KPC mice collected at the survival endpoint of the trametinib/SHP099 trial. Principal component analysis (PCA) of bulk tumor transcriptomes revealed that three of five vehicle-treated samples (samples 2, 4, and 5) clustered closely together and were therefore used as a reference (**Fig. S6e**). As expected, cell cycle regulation was re-induced in long-term dual-treated KPC-derived tissues, and heterogeneous remodeling of EMT-associated signatures, remained upregulated in 50% (4/8) of tissue samples (**Fig. 4c**).

Within category 3, sustained mTORC1 activation was observed after prolonged exposure. Most importantly, categories 4 and 5 showed a marked reduction in immune response and chemotaxis/migration pathways, respectively, in SHP099- or trametinib-only compared to dual therapy. Specifically, trametinib-treated KPC tissues exhibited strong attenuation of key immune pathways, including inflammatory signaling (6/8), IFN-α (5/8), IFN-γ (7/8), and CTLA-4 signaling (adj. p < 0.05), indicating suppression of antitumor immune activity. In contrast, dual MEK/SHP2 inhibition reprogrammed immune signaling rather than inducing uniform suppression, with only limited downregulation of IFN-α (1/8) and IFN-γ (2/8) pathways and partial pathway upregulation (2/8), consistent with a reduced immunosuppressive effect (**Fig. S6e; Fig. 4c**). These data supports the idea that adding SHP099 might partially attenuate trametinib-associated long term effects, in line with the reduced mMDSCs and gMDSCs infiltration shown by mIHC upon SHP099 addition.

CXCR3-associated chemokines, including CXCL9 and CXCL11, showed an overall decreasing trend, whereas CXCL10 showed a slight increasing tendency compared to monotherapies (**Fig. S7b**). In conclusion, long-term dual treatment suggests an adaptive resistance state characterized by diminished CXCR3-mediated chemokine signaling, which may potentially explain the decreased CD3^+^CD4^+^ T cell infiltration and the reaccumulation of M2-like macrophages observed in FC and mIHC over time.

### Single-cell profiling reveals immune cell functional alterations and suggests potential immunotherapeutic targets

#### I. Dynamic alterations in immune cell frequency, activation, and functional states

To achieve a more granular understanding of the PDAC tumor microenvironment and tumor cell state evolution under dual MEK/SHP2 inhibition in a real-time *in vivo* context, we then performed single-cell RNA sequencing. Fractions of CD45⁺ and CD45⁻ cells were isolated from 7 autochthonous KPC mice treated with vehicle, SHP099, Trametinib, or both at short- and long-term timepoints for separate sequencing. Uniform manifold approximation and projection (UMAP) and graph-based clustering identified 17 transcriptionally distinct clusters across CD45⁻ and CD45⁺ compartments, which were annotated using established marker genes from the literature (**Fig. 5a**).

**Fig. 5.**
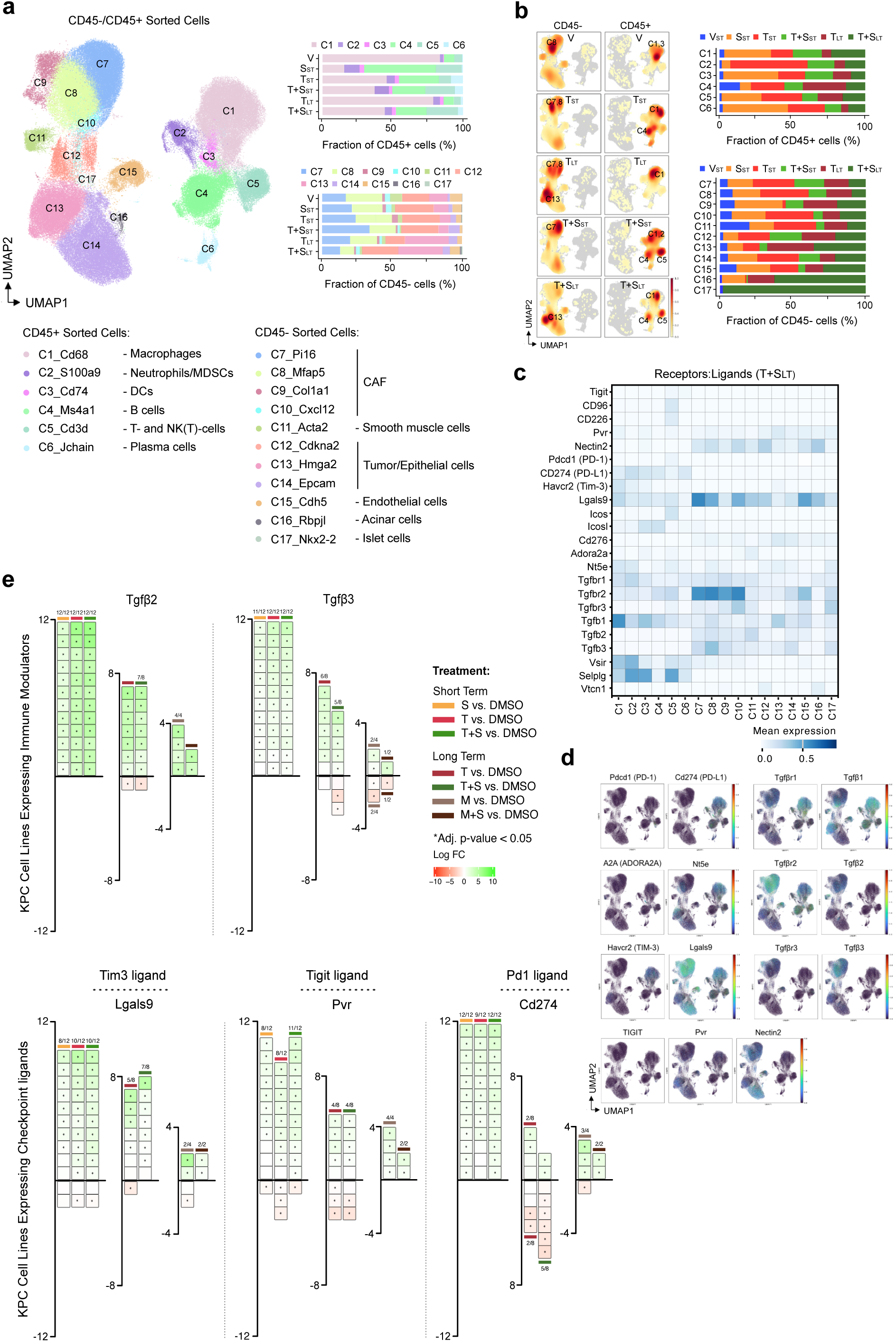
Single-cell profiling reveals immune cell functional alterations under dual MEK/SHP2 inhibition. **a.** UMAP plot of scRNA-seq data from CD45⁺ and CD45⁻ sorted cells showing 17 clusters, annotated based on marker gene expression. Bar plots show the distribution of each cluster across 6 treatment arms in CD45⁺ (top) and CD45⁻ (bottom) sorted cells. Treatment durations: ST, short term; LT, long term. Treatments: V, vehicle; S, SHP099; T, trametinib; T+S, trametinib + SHP099. **b.** Left panels show representative feature density overlays of sorted CD45⁺ and CD45⁻ cells, highlighting enrichment of selected clusters across treatment groups. Right panels show bar plots of the relative abundance of each treatment across clusters. **c**. Heatmap illustrating expression of checkpoint receptors, ligands and growth factors across clusters under T+S_LT_ treatment arm. **d.** Feature plots showing expression of selected checkpoint ligand-receptor pairs across UMAP clusters under T+S_LT_ treatment arm. **e.** Differentially expressed genes (DEGs) across KPC cell lines. Treatment durations: ST, short term; LT, long term. Treatments: V, vehicle; S, SHP099; T, trametinib; T+S, trametinib + SHP099; M, MRTX1133; M+S, MRTX1133 + SHP099. Color intensity indicates log fold change (logFC; red, downregulated; green, upregulated). Asterisks (*) denote significant differential expression (adjusted p < 0.05). Numbers above/below each group indicate the fraction of samples with significant changes per treatment arm.

Among the non-immune (CD45⁻) fraction, 11 dominant clusters corresponded to cancer-associated fibroblasts (CAFs) and epithelial tumor cells, alongside minor populations of smooth muscle cells, endothelial cells, acinar cells, and islet cells. In the immune (CD45⁺) compartment, 3 major populations were annotated as macrophages, T cells, and B cells, with smaller populations of neutrophils, dendritic cells, and plasma cells also detected. Cell type proportions varied across treatment groups, with T- and NK(T)-cells (C5_Cd3d) increasing after short-term SHP2 or dual MEK/SHP2 inhibition and remaining elevated under prolonged dual therapy, while macrophages (C1_Cd68) were consistently the most abundant immune population. In the CD45⁻ compartment, Hmga2⁺ tumor/epithelial cells decreased after short-term dual inhibition but rebounded with prolonged treatment, whereas Pi16⁺ and Mfap5⁺ CAF clusters were reduced under long-term therapy. Feature (density) UMAPs corroborate these findings, illustrating treatment-dependent redistribution of CD45⁺/CD45⁻ populations over time. The bar plots quantify the relative fraction of each cluster across treatment conditions (**Fig. 5a,b**).

We further analysed the (C5_Cd3d) cluster at higher resolution to define T- and NK(T)-cell states under MEK/SHP2 inhibition. Among the 13 identified clusters, we identified effector T-cells (C1, C2), helper T-cells (C3), naïve/memory T-cells (C4, C5), regulatory T-cells (C6, C7), NK and NK(T)-cells (C9, C10), γδ T-cells (C8), activated T-cells (C8), and additional low-abundance clusters (C12, C13). CD8⁺ T cell activation UMAP analyses further revealed treatment-dependent changes in activation states, alongside marked shifts in cluster composition across conditions. Vehicle-treated samples showed almost no evidence of CD8⁺ T cell activation. SHP099 treatment was associated with enrichment of naïve–memory CD4⁺ and CD8⁺ T cells, as well as Tregs and NK/NK(T)-cells. Short-term trametinib induced a clear early T cell activation signature, whereas long-term treatment resulted in loss of this activation profile, with persistence of a subset of early effector CD8⁺ T cells. Short-term combination treatment showed strong early effector CD8⁺ and γδ T cell activation. As expected, CD8⁺ T cell activation was not markedly enriched, as demonstrated by both FC and mIHC analyses. In contrast, long-term dual treatment was associated with a shift toward an effector-memory CD4⁺ reservoir (**Fig. S8a**).

Sub-clustering of macrophages (C1_Cd68) revealed distinct temporal enrichment of tumor-associated macrophages (TAMs) and condition-specific shifts in cluster composition. Vehicle-treated samples were enriched for two M2-like states, comprising homeostatic tissue-resident and activated remodeling macrophages. SHP099 treatment showed minimal evidence of TAM alteration. Short-term trametinib induced a pronounced tissue-resident M2-like signature, whereas prolonged exposure expanded C5, C6, and C8 subclusters, encompassing MHC-II–high antigen-presenting, hypoxia-associated inflammatory, and lipid-associated remodeling macrophages, respectively. Short-term combination therapy preferentially enriched MHC-II–high antigen-presenting macrophages, while long-term treatment favored monocyte-derived inflammatory populations (**Fig. S8b**). The reduced M2-like fraction observed by FC after short-term dual therapy therefore could reflect early reprogramming of resident M2-like cells into antigen-presenting and transitional TAM states, as supported by UMAP-defined C4 and C5 signatures.

#### II. MEK/SHP2 inhibition promotes T cell memory remodeling *in vitro*

Given the broad remodeling of the intratumoral T- and NK(T)-cell landscape under dual MEK/SHP2 inhibition, including expansion of naïve/memory T cell clusters, we performed a brief *in vitro* excursion to corroborate its impact on T cell memory differentiation, focusing on stem-like memory T cells (T_SCM_), which are critical for durable antitumor immunity due to their longevity, self-renewal, and multipotency. This is supported by previous findings that MEK inhibition alone has been shown to promote T_SCM_ generation with a naïve-like, highly proliferative capacity^32^.

First, we performed *ex vivo* assays using CD3/CD28 bead stimulation. Trametinib alone had minimal impact on T cell proliferation or viability, whereas combined MEK/SHP2 inhibition significantly reduced the number of viable CD8^+^ T cells. However, proliferation was not fully blocked, and pERK1/2 signaling persisted in both dividing and non-dividing cells, suggesting incomplete MAPK suppression. FC further showed that cytotoxic effector markers (Granzyme B, IFNγ, Perforin, CD107a) were largely maintained, or modestly increased, across inhibitor conditions in CD8⁺ T cells, suggesting preserved effector function despite pathway inhibition (**Fig. S9a**). To assess T_SCM_ cells in CD8⁺ T cell–moDC co-cultures, we examined T cell memory differentiation. The full experimental timeline is outlined in (**Fig. S9b**). Gating revealed a reduction in effector memory CD8⁺ subsets, alongside a preferential shift toward the central memory phenotype, least pronounced under dual treatment. Cytotoxic effector markers were maintained or modestly increased in T_SCM_ subsets following MEK/SHP2 inhibition, with the highest proportion of T_SCM_ cells observed in the SHP099 and combination arm (**Fig. S9c**). Collectively, the percentage of T_SCM_ cells was higher following SHP099 or combination treatment (**Fig. S9d**), potentially ascribing a beneficial impact of SHP2 inhibition on the T cell compartment. These observations, however, remain preliminary given the limited sample size and warrant further validation.

#### III. TIGIT and TIM-3 immune checkpoints and TGF-β as candidate immunotherapy targets

To find promising target candidates, we tracked the enrichment of each curated immune-related gene across all treatment arms from short- to long-term conditions and, within each cluster, highlighted genes exhibited three main dynamic patterns: unchanged across conditions (black), enriched in short-term (blue), or enriched in long-term (red) under dual treatment. Among these dynamically regulated genes, TGF-β1 showed sustained enrichment in C1 across all treatment arms and time points, while C13 displayed progressive enrichment under dual treatment. Other checkpoints, including Lgals9 and Pvr, also showed cluster-dependent patterns, being short-term enriched in some clusters but persistently enriched in others (**Fig. S8c**). Ligand–receptor pairs across clusters, highlighted Lgals9 (TIM-3 ligand), Nectin-2 (TIGIT ligand), and TGF-β ligands as prominent candidates enriched in CD45⁻/CD45^+^ populations (**Fig. 5c,d**).

We revisited transcriptomic profiles of our cell line panel under short- and long-term treatment to identify candidates for validation. KPC cell lines showed a robust short-term induction of TGF-β2/3 (12/12) and checkpoint ligands (Lgals9, Pvr, Cd276), which remained elevated in resistant states under both trametinib+SHP099 and MRTX1133+SHP099 (**Fig. 5e**). Long-term–treated human cell lines similarly upregulated TGF-β2 and multiple immune checkpoint ligands, including PD-L1, PD-L2, PVR (TIGIT ligand), 4-1BB, and to a lesser extent LGALS9 (**Fig. S7a**). In contrast, tissue samples showed an overall decrease in checkpoint ligands (PD-L2, PVR, CD276, NT5E) and LAG3, alongside partial upregulation of receptors such as TNFRSF25, TIGIT, and particularly TIM3 (3/8), likely reflecting changes in immune cell composition rather than tumor-intrinsic signaling (**Fig. S7b**). Taking into account the strong and consistent upregulation of TGF-β and TIGIT/TIM-3 checkpoint ligands in response to vertical RAS pathway inhibition across a large number of PDAC models, our findings suggest that these pathways could represent targets of value.

### Protein-level validation of TIGIT/TIM-3 checkpoints and TGFβ reveals early and persistent expression during tumor progression

To validate our transcriptomic findings at the protein level, we assessed TGF-β signaling, TIGIT ligands (PVR and NECTIN-2), and TIM-3 ligands (CEACAM1 and LGALS9) in murine and human cell lines, as well as treated KPC tissues.

Western blotting showed increased TGF-β across all human cell lines under short-term treatment, persisting in resistant cells (**Fig. S10a**). Classical lines showed similar short-term patterns, maintained in KPC-162 (**Fig. S10b**), whereas basal-like lines exhibited higher expression under both conditions; with MRTX1133+SHP099 effectively eliminating basal-like cells (**Fig. S10c**). Over time, TIM-3 ligand CEACAM1 increased under combination treatment in PANC-1 and MIA PaCa-II but decreased in most murine lines; LGALS9 protein expression could not be confirmed (data not shown; **Fig. S10a–c**). In contrast, the TIGIT ligand Nectin-2 showed persistent expression across nearly all human and basal murine lines, while PVR was also maintained, albeit at lower levels in murine cells. Overall, pSMAD2/3 showed variable changes (**Fig. S10a-c**).

Assessed by IHC, TIGIT and its ligand Nectin-2 (black) were the most abundantly expressed receptor–ligand pair across all conditions, while TIM-3 and its ligand Lgals9 (grey) showed consistently lower expression **(Fig. 6a)**. Normalized data show higher expression of TIGIT/Nectin-2 across all treatment arms. In contrast, TIM-3/Lgals9 remain detectable across all conditions, albeit at lower levels **(Fig. 6b)**. Bar plots display original, non-normalized cell counts (cells/mm²) to provide a direct view of absolute cell numbers (**Fig.S10d**). PD-1 was additionally assessed to confirm its comparatively minor role; although PD-1 and pSMAD2/3 signals increased across dual-treated groups, PD-1 contributed only minimally overall (**Fig. S10e,f**). Collectively, all proposed checkpoint candidates were detectable under short-term trametinib+SHP099 and persisted in long-term conditions. Their early presence and sustained expression in resistant states support their potential as combination immunotherapy targets already starting from treatment initiation.

**Fig. 6.**
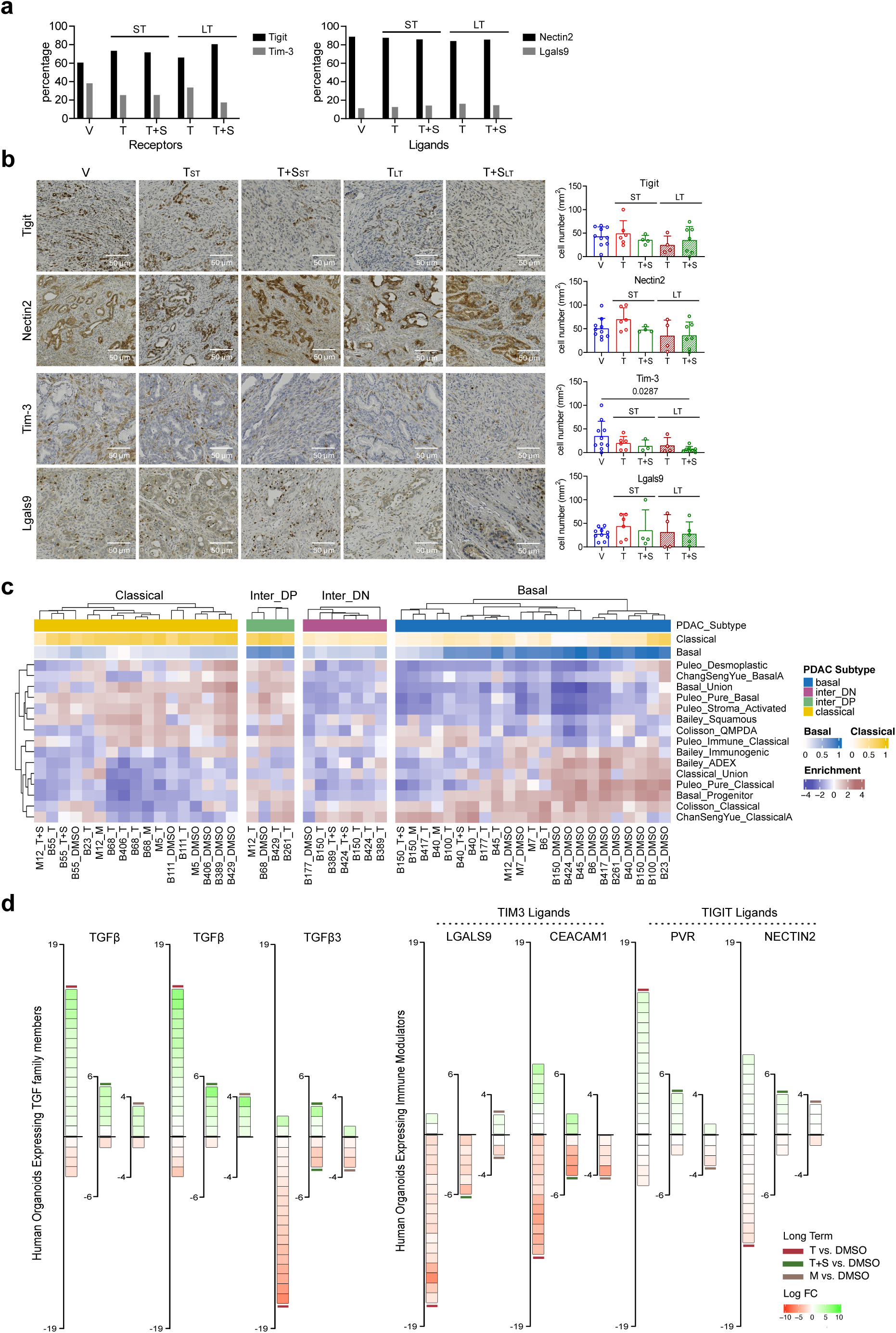
Proteomic and PDO-based validation support TGF-β and TIGIT/TIM-3 targeting as therapeutic strategies in PDAC. **a.** Bar plots show the percentage of receptor- and ligand-positive cells across all mice within each condition under short-term (ST) and long-term (LT) treatment. **b.** Representative IHC images of KPC tumors stained for TIGIT/Nectin-2 and TIM-3/LGALS9 under short-term (ST) and long-term (LT) treatment conditions. Treatments: T, trametinib; T+S, trametinib + SHP099. Scale bars, 50 μm. Dot plots (right) display individual tumors with mean ± SD. **c.** Subtype-resolved enrichment heatmap across Classical, Intermediate (DN: downregulated; DP: upregulated), and Basal states, highlighting therapy-induced transcriptomic reprogramming manifested as subtype transitions within the same patient derived organoid (PDO) line across resistant conditions. **d.** Differentially expressed genes (DEGs) across PDOs. Treatment durations: LT, long term. Treatments: T, trametinib; T+S, trametinib + SHP099; M, MRTX1133. Color intensity indicates log fold change (logFC; red, downregulated; green, upregulated).

### PDO-based validation support TIGIT/TIM-3 and TGF-β targeting as immunotherapeutic adjunct for vertical RAS pathway inhibition in human PDAC

For translational validation, we used 40 patient-derived PDAC organoids (PDOs) and treated over extended periods of time until they resumed stable proliferation. Overall, vertical RAS pathway inhibition was highly effective across the PDO lines, and resistance could only be established in fewer than 50% of models. In total, only 19 out of the 40 PDO lines developed resistance, with all 19/19 resistant to trametinib, 7/19 resistant to trametinib+SHP099, 4/19 resistant to MRTX1133, and 0/19 resistant to MRTX1133+SHP099 (**Fig. 6c**). ssGSEA was used to classify PDAC subtyping, revealing a spectrum ranging from classical to intermediate to basal-like subtypes.

Trametinib-resistant PDOs show, increased proliferation in basal-like PDOs, and suppressed immune response and migration/chemotaxis in intermediate and basal-like subtypes. Trametinib+SHP099- and MRTX1133-treated PDOs show a similar trend, alongside reduced immune programs in basal-like subtypes. Migration/Chemotaxis were largely absent across resistant settings (**Fig. S11a-c**). Overall, adaptive reprogramming across PDOs only partially recapitulated responses observed in 2D systems and the KPC *in vivo* model, highlighting a more heterogeneous and complex transcriptional landscape.

Resistant PDOs consistently downregulated CXCR3 ligands (CXCL9, CXCL10, CXCL11). In contrast, TGF-β1 and TGF-β2 were broadly upregulated following trametinib, trametinib+SHP099, and MRTX1133 treatment relative to DMSO, although without reaching statistical significance. Notably, TIGIT ligands showed pronounced upregulation relative to TIM-3 ligands and the PD-1 ligand (CD274), which were also present but in a less striking and less consistent manner (**Fig. S11d, Fig. 6d**). Together, the coordinated activation of TGF-β and TIGIT/TIM-3 signaling identifies these pathways as promising immunotherapeutic targets to complement vertical MAPK/SHP2 inhibition and potentially improve long-term PDAC control.

## Materials and Methods

### Mouse Models and Genetic Engineering

This study utilized the most common genetically engineered mouse models sufficient to induce invasive pancreatic ductal adenocarcinoma: Kras^tm1Tyj^: Kras^LSL-G12D/+^ + Trp53^tm1Brn^: Trp53^fl/fl^ + Ptf1a^tm1(Cre)Hnak^: Ptf1a^Cre-ex1/+^ (KPC), Kras^tm1Dsa^: Kras^FSF-G12D/+^; Trp53^tm1.1Dgk^: Trp53^frt/frt^; Ptf1a^tm1(flpo)Hcrd^: Ptf1a^Flpo/+^; Gt(ROSA)26Sor^tm3(CAG-Cre/ERT2)Dsa^: R26^FSF-CreERT/+^; Ptpn11^tm1Gsf^: Ptpn11^fl/fl^ (KPF-R26Cre-SHP2), both of which had a mixed genetic background. Genetic modifications were confirmed at weaning and post-mortem through polymerase chain reaction (PCR) and gel electrophoresis. Approval for animal experiments was granted by the local authority (Regierungspräsidium Freiburg, Germany; approval number G/19-137), ensuring strict compliance with institutional guidelines and relevant legal regulations. Additionally, the authors followed the “Animal Research: Reporting of in Vivo Experiments” (ARRIVE) guidelines. All mice were bred and housed under the same conditions, and all experiments were performed at the Medical Center - University of Freiburg.

### MRI Scanning

Magnetic resonance imaging (MRI) was performed using a 9.4 tesla small bore animal scanner (BioSpec 94/20, Bruker Biospin) and a dedicated mouse quadrature-resonator (Bruker) for in vivo mouse body imaging. Prior to scanning, animals were anesthetized at 2% isoflurane in oxygen, which was maintained at 1.5% - 3% isoflurane during imaging: Respiration was closely monitored to minimize motion artifacts that could interfere with the imaging of the pancreas. Core body temperature was maintained at 37°C using a temperature-controlled water heating system. Mice were screened weekly until a solid tumor mass was detected. KPC mice, however, continued to be monitored on a weekly basis to evaluate survival outcomes over time. The MRI protocol consisted of a localizer and a T2-weighted spin echo RARE (Rapid Acquisition with Relaxation Enhancement) sequence. The number of slices was adjusted to the measured volume (on average 25) to ensure complete coverage of the tumor.

### MR Tumor Volumetry

The total pancreatic volume was determined using MRI volumetry. For this method, a region of interest (ROI) encompassing the pancreas was manually delineated on each image slice using the web-based NORA software developed at the University Medical Center Freiburg, Germany. Volume was calculated by summing all voxel volumes within the ROI boundaries. Total volumes were derived from sets of contiguous images by summing the products of area measurements and slice thickness.

### Treatment Protocols and Study Design

KPC mice with tumors confirmed via MRI screening were treated via oral gavage. Over the course of either a short-term (2 weeks) or at survival point treatment regimen, the mice were randomly assigned to four treatment groups (at least n=13 per group) and assigned to one of the following treatments: 75 mg/kg SHP099 (Selleck Chemicals, S8278), 1 mg/kg. Trametinib (Selleck Chemicals, S2673), combination therapy with both drugs at the same doses, or vehicle control. Treatments were administered every other day. Trametinib and SHP099 were first dissolved in dimethyl sulfoxide (DMSO) and then diluted to prepare a final solution consisting of 0.4% DMSO, 0.8% Tween-10%, and 11.8% glucose (40% solution) in water.

### Flow Cytometry and Cytometry Analysis

To obtain a single-cell suspension, CD45+ cells were isolated from PDAC tumors in KPC-treated mice, as previously described^33^. Single-cell suspensions were incubated with Fc receptor blocking solution (BioLegend) for 5 minutes at room temperature and subsequently stained with fluorescently labeled antibodies in flow cytometry buffer (1x PBS, 5% FBS). For flow-cytometric analysis of lymphoid subpopulations, the following antibodies were used: PE/Cy5.5-TCRγ/δ (15-5711-81), BV785-LAG3 (125219), PE/Dazzle594-CD8a (100761), BV570-CD4 (100541), PerCP/Cy5.5-CD25 (101911), AF647-NK1.1 (108719), BV510-PD1 (135241), AF488-CD3 (152322), BV605-CD19 (115539), AF532-CD80 (NBP3-11976AF532), BV421-FOXP3 (126419), and PE/Cy7-T-bet (644823). For flow-cytometric analysis of myeloid subpopulations, the following antibodies were used: PE-Arg1 (12-3697-82), AF532-MHCII (58-5321-82), AF647-CD8a (100724), PerCP/Cy5.5-Ly6C (128011), PE/Cy7-CD11b (101215), BV421-CD86 (105031), BV510-Ly6G (127633), PE/Dazzle594-CD80 (104737), BV711-CD11c (117349), and APC/Cy7-F4/80 (123117). CD45 (Pacific Blue, 103126) was used for both lymphoid and myeloid panels. All antibodies were purchased from BioLegend, except for TCRγ/δ, Arg1, and MHCII from Invitrogen, and AF532-CD80 from Novus Biologicals. Intracellular staining was performed after fixation and permeabilization of extracellularly stained-cells using the IC-Fixation Buffer (Invitrogen). Flow cytometry was conducted using the Sony SP6800 Spectral Analyzer (Sony Biotechnology). Compensation, population analysis, and quantitation were carried out using FlowJo software (Version 10.10).

### Multiplex Immunohistofluorescence

Pancreatic tumor tissue samples were formalin-fixed and paraffin-embedded (FFPE). Sections of 4 μm thickness were cut from each FFPE block and mounted onto positively charged slides. Each tissue section underwent five sequential rounds of staining. Each round included incubation with a protein-blocking buffer (1% BSA/TBST, 5% normal goat serum) for 20 minutes at room temperature (RT), followed by application of the primary antibody overnight at 4°C. The sections were then incubated with the corresponding horseradish peroxidase-conjugated secondary polymer (EnVision K4003, Dako) for 60 minutes at RT. Tyramide signal amplification (TSA) was used to mediate covalent binding of Opal fluorophores to the target proteins. After five sequential staining rounds, sections were counterstained with DAPI (FP1490, Akoya Biosciences) and mounted with ProLong Diamond Antifade Mountant (P36961, Thermo Fisher Scientific). Lymphoid cell markers included CD3 (ab16669, Abcam), CD4 (ab183685, Abcam), CD8a (#98941, Cell Signaling Technology), Foxp3 (#126535, Cell Signaling Technology), and NK1.1 (#39197, Cell Signaling Technology). Myeloid cell markers included CD11b (ab133357, Abcam), CD103 (ab224202, Abcam), Ly6c (ab15627, Abcam), MHCII (ab180779, Abcam), CD11c (#97585, Cell Signaling Technology), F4/80 (#70076, Cell Signaling Technology), Ly6g (#87048, Cell Signaling Technology), and Arg1 (#93668, Cell Signaling Technology). Pan-cytokeratin (PanCK) was detected using antibody NB600-579SS (Novus Biologicals). TSA fluorophores used to bind various cell markers include: 4′6-diamidino-2-phenylindole (DAPI), Opal Polaris 480, Opal Polaris 520, Opal Polaris 570, Opal Polaris 620, Opal Polaris 690, and Opal Polaris 780 (Akoya Biosciences; Opal Polaris 7-Color Manual IHC Kit, NEL861001KT). All sections were scanned using the Phenocycler-Fusion imaging system (Akoya Biosciences). High-resolution images were processed with Phenocycler software (Akoya Biosciences) and subsequently analyzed using QuPath software (version 0.4.3), following previously described methodology^33^.

### Immunohistochemistry

Tissue sections were subjected to antigen retrieval using either EDTA buffer (pH 9.0) or citrate buffer (pH 6.0), depending on the target antigen. Primary antibodies were used as follows: pSmad2 (ab280888, Abcam; EDTA buffer pH 9.0; 1:2000), TIM-3 (ab241332, Abcam; EDTA buffer pH 9.0; 1:1500), PD-1 (ab309363, Abcam; EDTA buffer pH 9.0; 1:2000), TIGIT (ab233404, Abcam; citrate buffer pH 6.0; 1:500), Nectin-2 (ab135246, Abcam; citrate buffer pH 6.0; 1:500), and Galectin-9 (LGALS9; SAB4501751, Sigma-Aldrich; citrate buffer pH 6.0; 1:1000). Sections were incubated with primary antibodies overnight at 4 °C after blocking non-specific binding. Detection was performed using a secondary horseradish peroxidase-conjugated polymer (K4003, Dako), and signal was visualized using 3,3′-diaminobenzidine (DAB) (K3468, Dako). Sections were counterstained with hematoxylin (1.09249.0500, Merck), dehydrated, and mounted (ROTI®Histokitt II, T160.1, Carl Roth).

### Cell culture

Human PDAC cell lines, MIA PaCa-II, YAPC, and PANC-1, harboring KRAS^G12C^, KRAS^G12V^, and KRAS^G12D^, respectively, and their SHP2-knockout derivatives (generated via CRISPR-Cas9 targeting of PTPN11^34^) were used in this study. Cells were treated for 72 h to assess short-term responses and for at least 6 weeks to generate adaptive-resistant lines. Primary murine PDAC cell lines (F2453 and F2683), from the dual recombinase KPF model were formerly established, with Ptpn11 deletion induced by 2 μM 4-hydroxytamoxifen daily for six days^13^, while wild-type (WT) cells received vehicle, SHP099, trametinib, or both. Additional primary murine PDAC cells were obtained from freshly sacrificed KPC mice, cultured from ∼5 mm tumor fragments in RPMI-1640 with 10% fetal calf serum and 1% penicillin-streptomycin.

Following outgrowth, residual tissue was removed by trypsinization and cells were subcultured. For short-term experiments (48–72 h), cells were treated with 15 µM SHP099, 10 nM trametinib, a combination of SHP099 and trametinib, 10 µM MRTX1133, or a combination of MRTX1133 and SHP099. For long-term resistance assays, cells were continuously cultured from 6-10 weeks in the presence of the same drugs, with medium refreshed every three days and samples were collected at 90% confluence.

### Patient-derived *ex vivo* PDAC organoids

Primary human pancreatic ductal adenocarcinoma (PDAC) tissues were obtained from surgical resections at the Medical Center - University of Freiburg following written informed consent and approval by the institutional ethics committee (No. 126/17, 73/18), in accordance with the Declaration of Helsinki. PDAC diagnosis was confirmed by histopathological assessment. Fresh PDAC tissues were mechanically and enzymatically dissociated using collagenase, followed by ACK lysis, to generate single-cell suspensions. Cells were embedded in Matrigel and cultured as organoids in human growth medium based on established protocols^35^.

### T cell–DC co-culture

Peripheral blood mononuclear cells (PBMCs) were isolated from heparinized whole blood or buffy coat preparations obtained from healthy donors (Institut für Transfusionsmedizin und Gentherapie, Universitätsklinikum Freiburg) with informed consent and ethical approval (EK-Freiburg 21-1015). Monocytes were enriched from freshly isolated PBMCs using the Pan Monocyte Isolation Kit, human (Miltenyi Biotec, 130-096-537). Adherent monocytes were differentiated into monocyte-derived dendritic cells (moDCs) by culturing for six days in the same medium supplemented with GM-CSF (50 ng/mL; PeproTech, 300-03), IL-4 (20 ng/mL; PeproTech, 200-04) and a maturation cocktail containing TNF-α (10 ng/mL; PeproTech, 300-01A), IL-1β (10 ng/mL; PeproTech, 200-01B), IL-6 (1000 U/mL; PeproTech, 200-06), and PGE₂ (1 µg/mL; TOCRIS, 2296), without medium replacement. On day six, moDCs were harvested using Macrophage Detachment Solution DXF (Sigma-Aldrich, C-41330), counted, and resuspended for antigen loading prior to T cell co-culture^36,37^. For tumor neoantigen loading^36^, MIA PaCa-II tumor cells were lysed by repeated freeze–thaw cycles and sonication. The cleared lysate was added to moDC cultures at a 1:1 ratio and incubated for 48 h at 37 °C with 5% CO₂. FITC-Dextran (1 mg/mL; Sigma-Aldrich) was added to complete RPMI medium containing cytokines. Antigen-loaded moDCs were harvested as described above, centrifuged at 200 × g for 5 min at room temperature with brake 5, resuspended in 300 µL FACS buffer, and the FITC signal was measured by flow cytometry. T cells were isolated from PBMCs using the Pan T Cell Isolation Kit, human (Miltenyi Biotec, 130-096-535). T cells were labeled with carboxyfluorescein succinimidyl ester (CFSE; Biolegend, 423801). CFSE-labeled T cells were co-cultured with antigen-loaded moDCs at a 4:1 ratio in complete RPMI 1640 medium containing IL-2 (25 ng/mL final concentration; PeproTech, 200-02). Inhibitors, including SHP099, trametinib, or the combination of both, were added at the start of the co-culture. Cells were incubated for seven days at 37 °C with 5% CO₂, and culture medium was refreshed every 2–3 days. At the end of the assay, T cells were harvested and stained for flow cytometry analysis.

### T cell proliferation assay

For proliferation^32^, CFSE-labeled or unlabeled human T cells were cultured with either moDCs (as described above) and recombinant human IL-2 (25 ng/mL final concentration; PeproTech, 200-02) in complete RPMI 1640 medium supplemented with 10% heat-inactivated FBS and 100 U/mL penicillin/100 µg/mL streptomycin for seven days. Culture medium was refreshed every 2–3 days. Inhibitors (SHP099, trametinib, or the combination) were added where indicated. After seven days, T cells were harvested and analyzed by flow cytometry for proliferation, activation, exhaustion, cytotoxicity, and memory markers.

### Western blotting

Cells were washed with ice-cold PBS and lysed in RIPA buffer supplemented with protease inhibitor (HALT™ Protease Inhibitor Cocktail, Thermo Fisher Scientific, 78429) and phosphatase inhibitor (HALT™ Protease Inhibitor Cocktail, Thermo Fisher Scientific, 78420). Lysates were incubated on ice for 10 minutes, centrifuged at 16,000 × g for 5 minutes at 4°C, and supernatants were collected. Protein concentrations were measured using the BCA assay (Pierce BCA Protein Assay Kit, Thermo Fisher Scientific, 23225), and equal amounts were mixed with 1x Laemmli buffer containing β-mercaptoethanol, then denatured at 95°C for 15 minutes. Proteins were resolved on 10% SDS-PAGE, transferred to nitrocellulose membranes, and detected with the following antibodies: TGF-β (3711S), CEACAM1 (14771T), Nectin-2 (ab135246), PVR (ab103630), SMAD2/3 (8685T), pSMAD2/SMAD3 (8828T), pSHP2 (3751S), SHP2 (3397S), pSTAT3 (9145S), STAT3 (9139S), pAKT (13038T), AKT (4691S), pERK (4370S), ERK (4695S), MEK (8727S), pMEK (9154S), KRASG12D (53270S), E-Cadherin (3195S), N-Cadherin (13116S), GATA-6 (5851S), SLUG (9585S), SNAIL (3879S) and β-actin (4970S). β-actin was used at 1:5000 (Sigma-Aldrich), and all other antibodies were used at 1:1000; Nectin-2 and PVR were purchased from Abcam, and the remaining antibodies were obtained from Cell Signaling Technology. Signals were visualized with a ChemiDoc imaging system (Bio-Rad).

### Transcriptomics (RNA extractions, sequencing, and data analysis)

Total RNA was extracted from approximately 90% confluent cultured cells (human and murine PDAC cell lines) or ∼100 mg of frozen KPC pancreatic tumor tissue using the RNeasy® Plus Mini Kit (QIAGEN, 74136). RNA concentration was measured with a NanoDrop spectrophotometer, and RNA quality was assessed using an Agilent Bioanalyzer. Only RNA samples with an RNA integrity number (RIN) > 7.0 were included in the expression analysis. Sequencing of all RNA samples was performed at DKFZ using the Illumina NovaSeq™ 6000 sequencing system. Sequencing reads were converted to FASTQ format, quality-filtered, and mapped to the appropriate reference genomes (GRCm39 for mouse and GRCh38 for human) using the STAR aligner (version 2.7.10a)^38,39^. Downstream analysis was performed in R. Differentially expressed genes (DEGs) were analyzed using the limma package, while gene set enrichment analysis (GSEA) was conducted with the fgsea and MSigDB collections^40,41^. PDAC subtyping was perfomed by calculating a ssGSEA score based on Basal and Classical signatures, as described in Hafner et al^34^.

### Single-cell RNA Sequencing and Data Analysis

Single-cell RNA sequencing was performed using the GEXSCOPE® Single Cell RNA Library Kit (Singleron Biotechnologies). Sequencing data were processed with CeleScope (Singleron Biotechnologies, version 1.14.0) and aligned to the mouse reference genome Mus_musculus_ensembl_92 (Ensembl). Gene count matrices were further analyzed using Scanpy (version 1.9.1)^42^. Quality control was applied to retain only high-quality cells, defined by >200 detected genes, between 500 and 20,000 unique molecular identifiers (UMIs), and <15% mitochondrial gene content. The filtered data were normalized to total expression, scaled to the median UMI count across all cells, and log-transformed^43^. Principal component analysis (PCA) was performed without prior feature selection, utilizing all detected genes for dimensionality reduction and the data were subsequently visualized via UMAP^44,45^. Clustering was conducted using a range of resolutions (0.1–10) to identify a sufficient number of clusters, guided by the expression of known cell type markers. Differentially expressed genes (DEGs) were identified using the Wilcoxon rank-sum test, with significance thresholds set at an adjusted p-value ≤ 0.05 and log fold change ≥ 1. Gene set enrichment scores were calculated via Univariate Linear Model (ULM) as part of the decoupleR package^46^.

### Proteomics

Proteomic sample preparation, TMT labeling, fractionation was performed. 800 ng of peptides were analyzed using a Q-Exactive Plus mass spectrometer coupled to an EASY-nLC 1000 UHPLC system as described previously for LC-MS/MS. Data processing was conducted using MaxQuant v1.6.14.0 with the Human-EBI database (January 9, 2020 release)^34^.

### Cytokine Array

Primary murine SHP2 wild-type KPF tumor cell lines, F2453 & F2683, were cultured and treated with either SHP099, trametinib, combination, or a vehicle control for 48 hours. The tissue lysates were then analyzed using a Mouse XL Cytokine Array Kit (R&D Systems, ARY028) according to the manufacturer’s instructions. The levels of cytokines/chemokines were measured by determining the mean spot pixel intensity using ImageJ version 1.53a.

### Statistical Analysis

Statistical analyses were performed using GraphPad PRISM (Version 9.4.0). For experiments involving a single variable across multiple conditions, we used ordinary one-way ANOVA with multiple comparisons. A significance threshold of (P < 0.05) was applied to determine statistically significant differences between groups or samples. Survival data were assessed using Kaplan–Meier curves. High-dimensional omics data were analyzed in R Studio using the specified R packages^47^.

### Language Editing

Grammatical and stylistic improvements were carried out with the aid of an AI-based language model (ChatGPT). The model was used exclusively for linguistic refinement, and it did not contribute to data analysis, scientific interpretation, or the development of study content.

## Data availability

Transcriptomic and single-cell RNA-sequencing datasets generated in this study are publicly available. Bulk RNA-seq data can be accessed at [https://zenodo.org/records/16631280], and single-cell RNA-seq data can be accessed at [https://zenodo.org/records/16636353].

## Acknowledgments

We thank Tilmann Brummer and Nathalie Köhler for their valuable input and continued support as members of the thesis advisory committee (TAC).

- We thank the Core Facility Animal Imaging Research Center (AMIR), Department of Radiology – Medical Physics of the University Hospital Freiburg for support in acquisition (i.a. and analysis) of the data.
- We thank the Lighthouse Core Facility of the Medical Faculty, University of Freiburg (Project Numbers 2023/A2-Fol; 2021/B3-Fol), the DKTK, and the DFG (Project Number 450392965) for their support.
- We thank the Proteomics Platform – Core Facility (ProtCF) headed by Prof. Dr. Oliver Schilling (2021/A3-Sch; 2023/A3-Sch).

## Grant support and assistance

This study was funded by the German Cancer Aid, Deutsche Krebshilfe, Project ID 70113697 (to D.A.R.) and by the German Research Foundation, Deutsche Forschungsgemeinschaft (DFG), CRC1479 (Project ID 441891347, P17 to D.A.R., S01 to M.B, S02 to W.R.). A. Y. A. was supported by the **German Academic Exchange Service,** Deutscher Akademischer Austauschdienst (DAAD) and the ,Fill-in-the-Gap’ scholarship of the Faculty of Medicine, University of Freiburg. M.B. is supported by the DFG, within CRC1160 (Project ID 256073931, Z02), CRC/TRR167 (Project ID 259373024, Z01), CRC1453 (Project ID 431984000, S01), TRR 359 (Project ID 491676693, Z01), TRR 353 (Project ID 471011418-SP02), and FOR 5476 UcarE (Project ID 493802833, P07). M.B. also acknowledges funding from the German Federal Ministry of Education and Research (BMBF) within the Medical Informatics Funding Scheme, PM4Onco–FKZ 01ZZ2322A. G.A. is supported by the BMBF via EkoEstMed–FKZ 01ZZ2015. J.D.L. is funded by the Research Commission of the Faculty of Medicine, University of Freiburg.

## Contributions

The study was initially designed and conceived by D.A.R., with him also supervising the project and providing overall guidance. A.Y.A. performed the majority of experiments, data analysis and generated most of the figures. A.Y.A. also wrote the manuscript draft. X.C., P.H., S.K., H.J., and S.B. contributed to mouse experiments, sample processing, and cell lines treatment. Transcriptomic and bioinformatic analyses were performed by T.D. and G.A. under the supervision of M.B.. S.H., S.M., and K.M. contributed to IHC, multiplex staining, and patient-derived organoid work. Additional experiments and analyses were carried out by T.Y.A., M.A. and M.S.. D.V.E. and W.R. facilitated the mouse MRI studies. J.D.L. provided clinical input and critical feedback. D.A.R. secured funding. All authors reviewed and approved the final version of the manuscript.

## Discussion

Several studies have demonstrated the efficacy of SHP2-based combination therapies in KRAS-driven cancers, including PDAC^13,14,18,48^. Therapeutic effects of SHP2-based combinations extend beyond direct actions on cancer cells to modulation of immune components within the TME^18,49,50^. Nevertheless, potential adverse effects of SHP2 inhibition on tumor-associated immune cells may limit the potential of SHP2 inhibitors alone or in combination^49^. The balance between therapeutic benefit and adverse effects, and particularly the durability of these effects over time, remains incompletely understood.

Here, we find that dual inhibition transiently remodels the TME toward a likely beneficial, more pro-inflammatory state, characterized by increased T cell infiltration and a reduction in M2-like macrophages, alongside delayed tumor progression, and reduced proliferative signaling. However, these effects are also associated by a decrease in mDCs and a concomitant expansion of mMDSCs, indicative of a mixed immunological response. These changes are associated with tumor cell-intrinsic upregulation of CXCR3 ligands and TGF-β, as well as increased expression of checkpoint ligands, especially for TIGIT and TIM-3, across molecular subtypes and species. Over time, with the transition to an adaptive resistant tumor, this short-lived partial immune sensitization is lost, and immune-suppressive features prevail. M2-like macrophages re-accumulate and dendritic cell maturation remains impaired. In parallel, TGF-β expression persists, and TIGIT and TIM-3 ligand expression is further enhanced.

Our FC, mIHC, and transcriptomic analyses reveal that dual MEK/SHP2 inhibition induces an IFNγ-associated chemokine program, including CXCL10 and CXCL11. This is consistent with prior studies showing that MAPK-pathway inhibition induces de novo transcription of IRF1 and RIG-I (DDX58), which, upon IVT4 binding, drives TBK1/IRF3-independent inflammatory reprogramming of MAPK-targeted therapies^51^, and can up-regulate IFN signaling via MYC inhibition^52^. In our setting, however, this IFNγ-associated chemokine program fails to elicit a robust cytotoxic CD8⁺ T cell response. Instead, it coincides with selective enrichment of CD4⁺ T cells, indicating immune priming and remodeling of the tumor microenvironment rather than effective execution of anti-tumor immunity. Although CXCR3 ligands are classically critical to effector T cell recruitment^23^, their induction in PDAC does not necessarily translate into functional CD8⁺ responses. This likely reflects persistent stromal barriers or T cell dysfunction rather than insufficient chemokine signaling. Notably, the low abundance of cross-presenting dendritic cells, i.e. type 1 conventional DCs like CD11c⁺MHCII⁺CD8a⁺ and CD11c⁺MHCII⁺CD103⁺ DCs, likely limits MHC-I-mediated priming of naïve CD8⁺ T cells, potentially explaining why IFNγ-driven chemokine induction does not translate into CD8⁺ T cell enrichment. However, dual inhibition seems to promote the expansion of a memory T cell reservoir, particularly stem-like memory (T_SCM_) CD8+ T cells, a self-renewing population with preserved effector potential, previously shown to be induced by MEK inhibition alone^32^.

Dual MEK/SHP2 inhibition not only enhanced T cell infiltration, particularly CD4⁺ T cells, but also significantly reduced M2-like macrophages, indicating early repolarizastion shift of resident M2-like cells into antigen-presenting and transitional TAM states, as supported by single cell sequencing. The resulting shift improved the M1/M2 macrophage ratio, highlighting early myeloid remodeling. Transcriptional profiling reinforced this observation, revealing activation of innate immune programs, including myeloid leukocyte migration and chemotaxis, hallmarks of an immune-sensitized microenvironment. Transcriptomics profiling of human PDAC cell lines, as confirmed previously^49^, showed persistent upregulation of CXCL1 and CXCL5, whose protein products signal via CXCR2, a key chemotaxis receptor for MDSCs^53^, as well as CCL5, which may contribute to M2 macrophage depletion via NF-κB activation and promotion of M1 polarization^54^. This signature coincided with the emergence of mMDSCs following short-term dual inhibition and the re-emergence of M2-like macrophages during prolonged treatment, as detected by FC.

Prior studies have revealed immunological limitations of MAPK pathway inhibition and proposed complementary strategies to improve durable anti-tumor immunity. One study showed that pulsatile MEK inhibition preserves T cell activation and proliferation, delays resistance, and, when combined with CTLA-4 blockade, improves anti-tumor efficacy and survival in KRAS-mutant NSCLC^55^. Another study reported that some tumors become immunoresistant, particularly after acquiring resistance to MAPK inhibition, due to enhanced transcriptional output driven by reactivation of the MAPK pathway, and that restoring functional CD103⁺ DCs can resensitize these cross-resistant tumors to immunotherapy^56^. The same group also suggested that effective anti-tumor immunity relies on intratumoral CD8⁺ T cell priming by inflammatory monocytes with CXCL9⁺/CXCL10⁺ macrophage-like features. However, oncogenic MAPK signaling in tumor cells can disrupt this process through regulation of PGE2 and type I interferon responses, facilitating immune evasion. They proposed a mechanism-based combination of immunotherapy with COX-2i, 5-AZA, and FLT3L, suppressing PGE2-mediated immune evasion, restoring IFN-I responses, and enhancing cDC-mediated T cell priming, respectively^57^.

Intriguingly, our scRNA-seq data, western blotting, and IHC staining revealed upregulation of TIGIT axis (NECTIN2, PVR), TIM-3 axis (CEACAM1, LGALS9), and TGF-β isoforms, but not checkpoint receptors including PD1, CTLA4, or LAG3. These findings suggest that alternative checkpoints could be of value in PDAC, which largely does not respond to conventional PD1/PD-L1 blockade^58^. Co-targeting the CD155/TIGIT axis alongside PD-1 blockade and CD40 agonist–based therapy has been shown to bear potential for eliciting anti-tumor responses^59^. Others reported an overall 75% disease control rate upon the addition of the TGF-β receptor type I small-molecule inhibitor LY3200882 to gemcitabine and nab-paclitaxel in treatment-naive patients with advanced pancreatic cancer^60^. Positioning these pathways as compelling targets, suggests that combined TIGIT/TIM-3 ICB blockade and TGF-β antagonism may have the potential to improve long-term therapeutic response and help overcome resistance in the context of SHP2-based vertical MAPK pathway inhibition in PDAC^50,59,60^.

Recent advances in direct RAS inhibition have shifted the therapeutic landscape, rendering MEK or ERK inhibition comparatively less favorable. Nonetheless, SHP2 inhibition remains a compelling combinatorial strategy alongside RAS inhibitors due to its upstream position in RTK-RAS signaling and its immunomodulatory functions. A number of studies have investigated immunotherapy combinations as potential therapeutic partners for direct RAS inhibition. One group reported that especially αCTLA-4 blockade was substantially effective (without added value of αPD-1) in sustaining MRTX1133-mediated tumor control and CD8⁺ T-cell infiltration, partly via FAS-FASL engagement^61^. Others demonstrated how the RAS(ON) multi-selective inhibitor RMC-7977 achieved the strongest anti-tumor responses when combined with αCTLA-4, αPD-1, and CD40 agonism, with CD40 emerging as the dominant contributor, likely by boosting T cell priming via CD103⁺ DCs^62^. In a related approach, combining MRTX1133 with CXCR1/2 inhibitor, αLAG3, and agonistic α41BB synergistically reprogrammed the PDAC tumor microenvironment into a highly immunogenic state by depleting gMDSCs, promoting M1-like antigen-presenting macrophages, expanding CD103⁺ cross-presenting cDC1s, and enhancing CD8⁺ T-cell priming and infiltration^63^. Together, these reports support a premise that future immunotherapeutic strategies likely need to employ a multi-pronged ‘cocktail’ approach, when intended to complement RAS pathway targeted therapies.

Our data suggest that SHP2-based vertical RAS pathway inhibition blockade could not only delay adaptive resistance but also create an opportunity for enhancing anti-tumor immunity specifically by targeting the TIGIT and TIM-3 checkpoint axes and TGF-β signaling, thereby potentially improving the durability of tumor control.

