## Supplementary Figures for "Immunomodulatory impact of *PTPN11*/SHP2-based vertical RAS-MAPK pathway inhibition in pancreatic cancer"

**Supplementary Figure Legends:**

**Fig. S1 | Dual MEK/SHP2 inhibition alters lymphoid and myeloid cell populations in PDAC. a.** Flow cytometry–based quantification of lymphoid and myeloid cell populations. Data are shown for each treatment group at short-term (2 weeks) and survival endpoints. Statistical analysis was performed using one-way ANOVA across the cohort; bars represent mean ± SD. Significance is indicated as $p < 0.05$ and $p < 0.005$. Frequencies are shown relative to the $CD45^+$ population. **b.** Gating strategy of lymphoid and myeloid populations. Live singlets were gated on $CD45^+$ leukocytes.

**Fig. S2 | mIHC reveals the broader immune context of targeting MEK/SHP2 in PDAC.** Collated data from autochthonous KPC PDAC tumors assessed by mIHC (cells/mm²), including **a,** T cells and subsets (Tregs and NKT cells), myeloid cells and dendritic-like cells. Statistical comparisons across the cohort were performed using one-way ANOVA; significant differences are indicated ($p < 0.05$). **b.** Single-channel images of FFPE OPAL-stained sections acquired using Akoya imaging systems and processed with Inform and QuPath.

**Fig. S3 | mIHC validates macrophage state shifts following trametinib and GS493 treatment. a.** FFPE PDAC tumor sections from the four treatment arms were stained using OPAL multiplex immunofluorescence across three immune panels: (i) “Lymphoid cell panel” (PanCK, CD3, CD4, CD8, NK1.1, FOXP3), (ii) “Myeloid cell panel” (CD11b, F4/80, Arg1, Ly6G, Ly6C), and (iii) “Dendritic cell panel” (CD11c, MHCII, CD103, CD8, CD11b). Whole sections were imaged at 20× magnification using the Akoya PhenoImager HT system. Immune context was quantified across panels and plotted in GraphPad Prism as cell density (cells/mm²) for each treatment arm at short-term and survival time points. **b.** Collated data are shown assessed by mIHC (cells/mm²), of T cells ($CD3^+$), myeloid cells ($CD11b^+$), and dendritic cells ($CD11c^+$). Statistical analysis was performed across the cohort using one-way ANOVA; bars represent mean ± SD, and significant differences are indicated as $p < 0.05$, $p < 0.005$, and $p < 0.0005$. Treatment durations: LT, long term. Treatments: V, vehicle; G, GS493; T, trametinib; T+G, trametinib + GS493.

**Fig. S4 | Proteomic screening for immune-related changes in murine KPF *in vitro* a,b** Western blot analysis of KPF F2683 and F2453 cells SHP2 WT treated with S, SHP099; T, trametinib; T+S, trametinib + SHP099 and SHP2 KO treated with T, trametinib, for 48 h. β-actin served as loading control. **c,d** Secretome profiling of KPF F2683 and F2453 cells after 48 h treatment; radar plots show chemokine signatures across conditions based on cytokine/chemokine arrays. **e,f** Quantification of secreted chemokines and interleukins in KPF F2683 and F2453 cells under the indicated conditions.

**Fig. S5 | Proteomic screening for immune-related changes in human PDAC cell lines *in vitro*. a.** Proteomic profiling of human PDAC cell lines demonstrates altered protein abundance following 72 hours of treatment. Displayed are proteins with Log2 fold change < -0.5 or > 0.5, and adjusted P-values < 0.05. **b.** GSEA comparing MIA PaCa-2 and YAPC cells treated with T or T+S versus DMSO using HALLMARK gene sets. **c**. Heatmap showing enrichment scores for published PDAC subtype signatures across human pancreatic cancer cell lines, annotated with morphological classification (basal-like, classical, intermediate), Western blot markers, and calculated Basal/Classical subtype scores. **d.** Gene set enrichment analysis (GSEA) of significantly enriched Hallmark, GO_BP, KEGG (adjusted $p < 0.05$), and BIOCARTA ($p < 0.05$) pathways, grouped into five functional categories: cell cycle regulation, tissue remodeling, oncogenic signaling, immune response, and migration/chemotaxis. Normalized enrichment scores (NES) are shown (blue, downregulated; red, upregulated). Sample type, database, treatment, and duration are indicated below. Treatment durations: ST, short term; LT, long term. Treatments: S, SHP099; T, trametinib, T+S, trametinib+SHP099. The analysis includes human cell lines under ST and LT conditions, stratified by PDAC subtype (Classical, Intermediate, Basal).

**Fig. S6 | Transcriptional profiling reveals distinct biological and immune-driven programs associated with treatment response and molecular subtypes in PDAC. a,b** Heatmap showing enrichment scores for published PDAC subtype signatures in untreated murine KPC-derived PDAC cell lines. Samples were stratified into Basal, Intermediate, and Classical groups based on integrated subtype signatures. Protein-level validation of subtype-associated markers by Western blot; β-actin served as the loading control. Protein lysates were collected from passages 5–10. **c-e** Gene set enrichment analysis (GSEA) of significantly enriched Hallmark, GO_BP, KEGG (adjusted $p < 0.05$), and BIOCARTA ($p < 0.05$) pathways, grouped into five functional categories: cell cycle regulation, tissue remodeling, oncogenic signaling, immune response, and migration/chemotaxis. Normalized enrichment scores (NES) are shown (blue, downregulated; red, upregulated). Sample type, database, treatment, and duration are indicated below. Treatment durations: ST, short term; LT, long term. Treatments: S, SHP099; T, trametinib. **c,d** KPC-derived cell lines treated with S or T; PDAC subtypes (Classical, Intermediate, Basal) indicated above. **e.** KPC bulk tumors treated with S or T. Principal component analysis (PCA) of transcriptomes shows that vehicle samples 2, 4, and 5 cluster closely together and and used as a common reference group.

**Fig. S7 | Treatment-induced DEGs in human PDAC cell Lines and KPC bulk tissues. a,b** Differentially expressed genes (DEGs) in human PDAC cell lines and KPC bulk tissues under short-term and long-term treatment conditions. Adjusted p-value < 0.05; green, upregulated; red, downregulated.

**Fig. S8 | Single-Cell profiling reveals immune cell functional alterations under dual MEK/SHP2 inhibition. a.** UMAP plot of scRNA-seq data from T- and NK(T)-cells showing 13 sub-clusters, annotated based on marker gene expression. Bar plots show the distribution of each cluster across 6 treatment arms (Top), and the relative abundance of each treatment across clusters (Bottom). $CD8^+$ T cell activation UMAP analyses present treatment-dependent changes in $CD8^+$ T cell activation states, and cluster composition across conditions. **b.** UMAP plot of scRNA-seq data from macrophages showing 9 sub-clusters, annotated based on marker gene expression. Bar plots show the distribution of each cluster across 6 treatment arms (Top), and the relative abundance of each treatment across clusters (Bottom). TAM activation UMAP analyses present treatment-dependent changes in TAM activation states, and cluster composition across conditions. **c.** Hierarchical heatmap and cluster-specific gene modules. colors indicate temporal treatment response: unchanged across conditions (black), enriched in short-term (blue), or enriched in long-term (red) under dual treatment.

**Fig. S9 | MEK/SHP2 inhibition promotes T cell memory remodeling *in vitro.* a.** CD3/CD28 bead–stimulated $CD8^+$ T cells analyzed for proliferation (CFSE dilution), pERK1/2 signaling in dividing and non-dividing populations, and effector function (Granzyme B, IFN-γ, Perforin, CD107a). **b.** Schematic timeline of the $CD8^+$ T cell–moDC co-culture experiment. **c.** Flow cytometry gating for $CD8^+$ T cell co-culture subsets (naïve, central memory [CM], effector memory [EM], and stem cell memory [$T_{SCM}$]) with functional marker assessment across subsets and treatment conditions. **d.** Quantification of $CD8^+$ T cell differentiation states showing the relative distribution of naïve, effector, and $T_{SCM}$ populations, (mean ± SD).

**Fig. S10 | Protein-level validation of TIGIT/TIM-3 checkpoints and TGFβ reveals early and persistent expression during tumor progression. a.** Immunoblot analysis of three human PDAC cell lines (YAPC, PANC-1, MIA PACA II); β-actin served as loading control. Molecular weight markers (kDa) are indicated. **b,c.** Immunoblot analysis of primary KPC tumor–derived cell lines (classical: KPC 162, 215; basal: KPC 382, 495). **d.** Bar plots showing raw (non-normalized) cell counts (cells/$mm^2$), providing absolute quantification. **e.** Bar plots showing the percentage of PD1 within each condition under short-term (ST) and long-term (LT) treatment. **f.** Representative IHC images of KPC tumors stained for PD1 and pSMAD2/3; scale bars, 50 µm. Dot plots show individual tumors with mean ± SD (right). Treatment durations: ST, short term; LT, long term. Treatments: V, vehicle; S, SHP099; T, trametinib; T+S, trametinib + SHP099; M, MRTX1133; M+S, MRTX1133 + SHP099.

**Fig. S11 | Transcriptional profiling reveals distinct biological and immune-driven programs associated with treatment response and molecular subtypes in PDOs.** Gene set enrichment analysis (GSEA) of significantly enriched Hallmark, GO_BP, KEGG (adjusted

$p < 0.05$), and BIOCARTA ($p < 0.05$) pathways, grouped into five functional categories: cell cycle regulation, tissue remodeling, oncogenic signaling, immune response, and migration/chemotaxis. Normalized enrichment scores (NES) are shown (blue, downregulated; red, upregulated). Sample type, database, treatment, and duration are indicated below. Treatment durations: ST, short term; LT, long term. Treatments: T, trametinib; T+S, trametinib + SHP099; M, MRTX1133. PDAC subtypes: Classical, Intermediate (DN: downregulated; DP: upregulated), and Basal states. **a.** Patient derived organoids (PDOs) treated with T under LT conditions. **b.** PDOs treated with T+S under LT conditions. **c.** PDOs treated with M under LT conditions. **d.** DEGs across KPC cell lines. Color intensity indicates log fold change (logFC; red, downregulated; green, upregulated).

**Fig. S1**

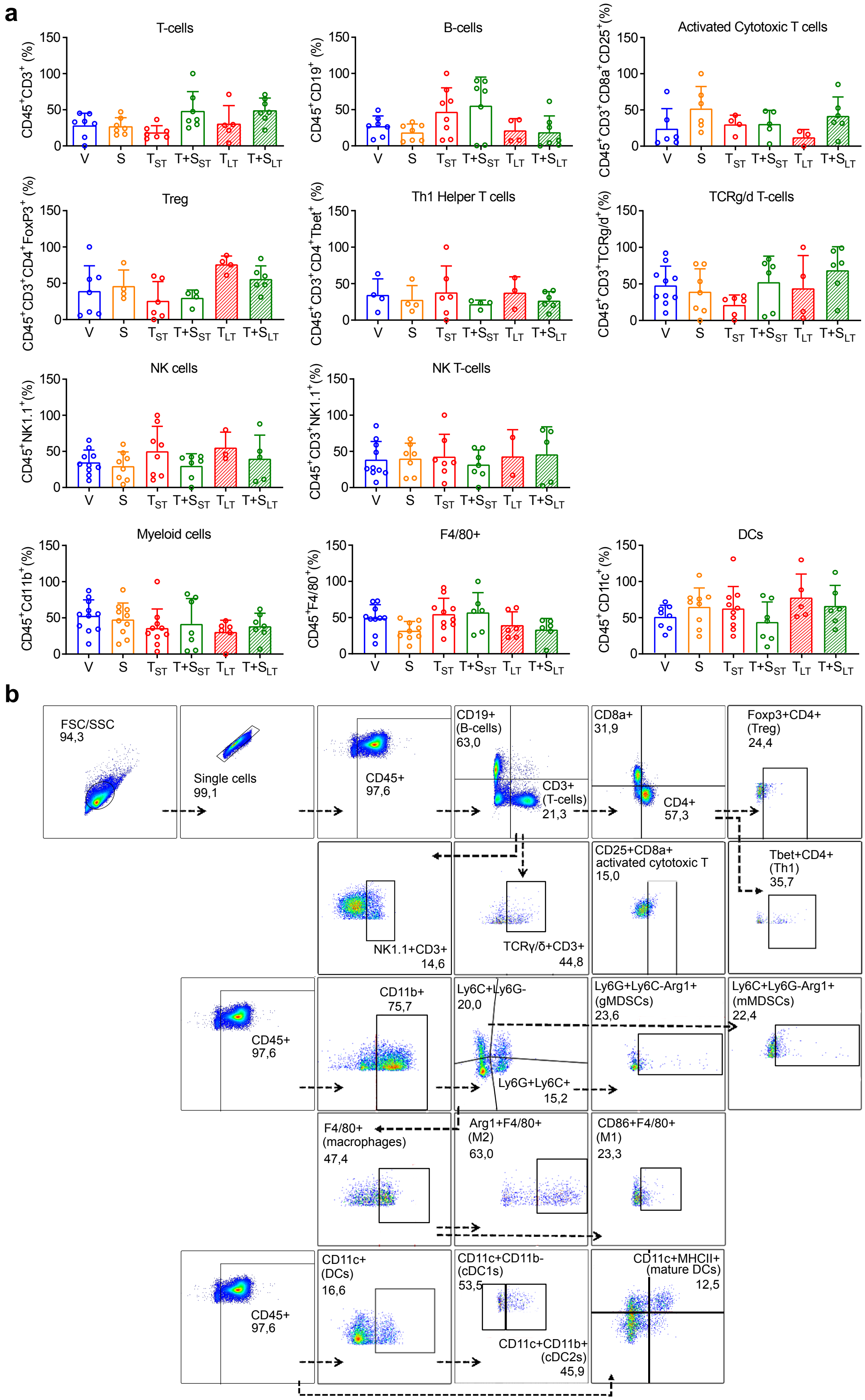
a
T-cells
CD45+CD3+ (%)
B-cells
CD45+CD19+ (%)
Activated Cytotoxic T cells
CD45+CD3+CD8a+CD25+(%)
Treg
CD45+CD3+CD4+FoxP3+ (%)
Th1 Helper T cells
CD45+CD3+CD4+Tbet+ (%)
TCRg/d T-cells
CD45+CD3+TCRg/d+(%)
NK cells
CD45+NK1.1+(%)
NK T-cells
CD45+CD3+NK1.1+(%)
Myeloid cells
CD45+Cd11b+(%)
F4/80+
CD45+F4/80+(%)
DCs
CD45+CD11c+(%)
0
50
100
150
V
S
TST
T+SST
TLT
T+SLT
b
FSC/SSC
94,3
Single cells
99,1
CD45+
97,6
CD19+
(B-cells)
63,0
CD3+
(T-cells)
21,3
CD8a+
31,9
CD4+
57,3
Foxp3+CD4+
(Treg)
24,4
NK1.1+CD3+
14,6
TCRγ/δ+CD3+
44,8
CD25+CD8a+
activated cytotoxic T
15,0
Tbet+CD4+
(Th1)
35,7
CD45+
97,6
CD11b+
75,7
Ly6C+Ly6G-
20,0
Ly6G+Ly6C+
15,2
Ly6G+Ly6C-Arg1+
(gMDSCs)
23,6
Ly6C+Ly6G-Arg1+
(mMDSCs)
22,4
F4/80+
(macrophages)
47,4
Arg1+F4/80+
(M2)
63,0
CD86+F4/80+
(M1)
23,3
CD45+
97,6
CD11c+
(DCs)
16,6
CD11c+CD11b-
(cDC1s)
53,5
CD11c+CD11b+
(cDC2s)
45,9
CD11c+MHCII+
(mature DCs)
12,5

#### Fig. S2

**a**

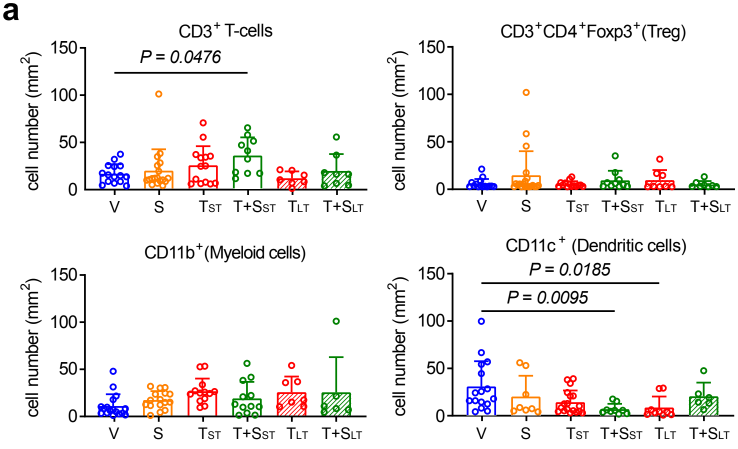


**b**

**Single Channel Images**

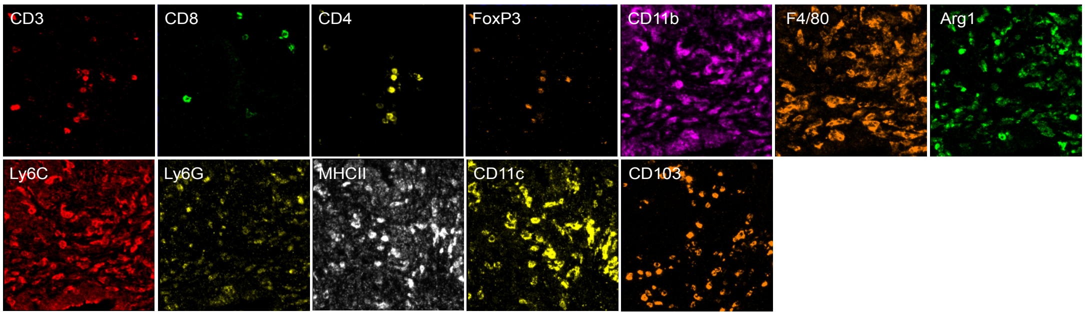

### Fig. S3

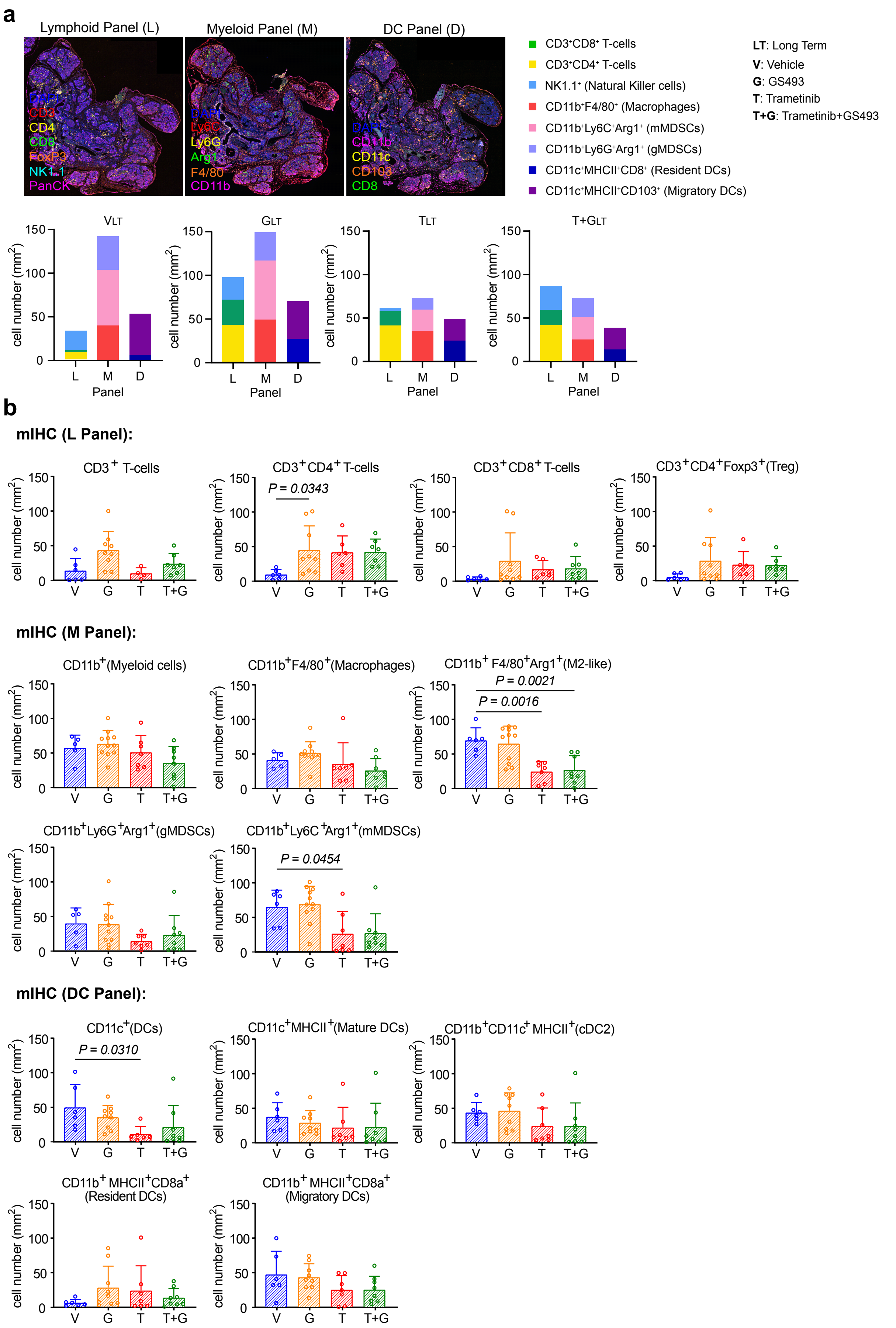
a
Lymphoid Panel (L)
Myeloid Panel (M)
DC Panel (D)
DAPI
CD3
CD4
CD8
FoxP3
NK1.1
PanCK
DAPI
Ly6C
Ly6G
Arg1
F4/80
CD11b
DAPI
CD11b
CD11c
CD103
CD8
CD3$^+$CD8$^+$ T-cells
CD3$^+$CD4$^+$ T-cells
NK1.1$^+$ (Natural Killer cells)
CD11b$^+$F4/80$^+$ (Macrophages)
CD11b$^+$Ly6C$^+$Arg1$^+$ (mMDSCs)
CD11b$^+$Ly6G$^+$Arg1$^+$ (gMDSCs)
CD11c$^+$MHCII$^+$CD8$^+$ (Resident DCs)
CD11c$^+$MHCII$^+$CD103$^+$ (Migratory DCs)
**LT**: Long Term
**V**: Vehicle
**G**: GS493
**T**: Trametinib
**T+G**: Trametinib+GS493
VLT
GLT
TLT
T+GLT
cell number (mm$^2$)
Panel
L
M
D
b
mIHC (L Panel):
CD3$^+$ T-cells
CD3$^+$CD4$^+$T-cells
CD3$^+$CD8$^+$T-cells
CD3$^+$CD4$^+$Foxp3$^+$(Treg)
P = 0.0343
mIHC (M Panel):
CD11b$^+$(Myeloid cells)
CD11b$^+$F4/80$^+$(Macrophages)
CD11b$^+$ F4/80$^+$Arg1$^+$(M2-like)
P = 0.0021
P = 0.0016
CD11b$^+$Ly6G$^+$Arg1$^+$(gMDSCs)
CD11b$^+$Ly6C$^+$Arg1$^+$(mMDSCs)
P = 0.0454
mIHC (DC Panel):
CD11c$^+$(DCs)
P = 0.0310
CD11c$^+$MHCII$^+$(Mature DCs)
CD11b$^+$CD11c$^+$ MHCII$^+$(cDC2)
CD11b$^+$ MHCII$^+$CD8a$^+$
(Resident DCs)
CD11b$^+$ MHCII$^+$CD8a$^+$
(Migratory DCs)
V
G
T
T+G
0
50
100
150

### Fig. S4

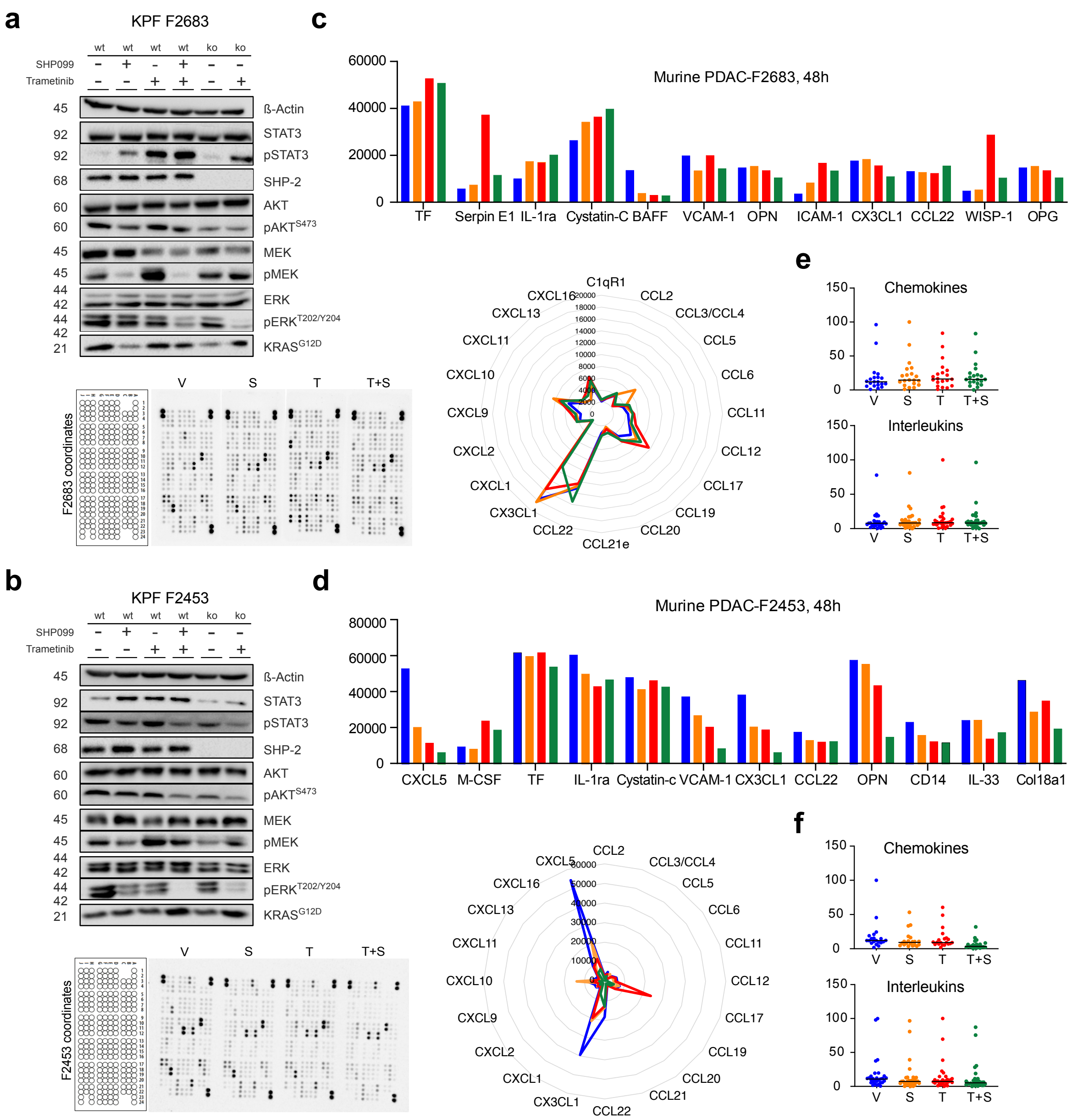

a
KPF F2683
wt wt wt wt ko ko
SHP099
Trametinib
ß-Actin
STAT3
pSTAT3
SHP-2
AKT
pAKTS473
MEK
pMEK
ERK
pERKT202/Y204
KRASG12D
F2683 coordinates
V S T T+S
b
KPF F2453
wt wt wt wt ko ko
SHP099
Trametinib
ß-Actin
STAT3
pSTAT3
SHP-2
AKT
pAKTS473
MEK
pMEK
ERK
pERKT202/Y204
KRASG12D
F2453 coordinates
V S T T+S
c
Murine PDAC-F2683, 48h
TF Serpin E1 IL-1ra Cystatin-C BAFF VCAM-1 OPN ICAM-1 CX3CL1 CCL22 WISP-1 OPG
C1qR1 CCL2 CCL3/CCL4 CCL5 CCL6 CCL11 CCL12 CCL17 CCL19 CCL20 CCL21e CCL22 CX3CL1 CXCL1 CXCL2 CXCL9 CXCL10 CXCL11 CXCL13 CXCL16
d
Murine PDAC-F2453, 48h
CXCL5 M-CSF TF IL-1ra Cystatin-c VCAM-1 CX3CL1 CCL22 OPN CD14 IL-33 Col18a1
CCL2 CCL3/CCL4 CCL5 CCL6 CCL11 CCL12 CCL17 CCL19 CCL20 CCL21 CCL22 CX3CL1 CXCL1 CXCL2 CXCL9 CXCL10 CXCL11 CXCL13 CXCL16 CXCL5
e
Chemokines
Interleukins
V S T T+S
f
Chemokines
Interleukins
V S T T+S

#### Fig. S5

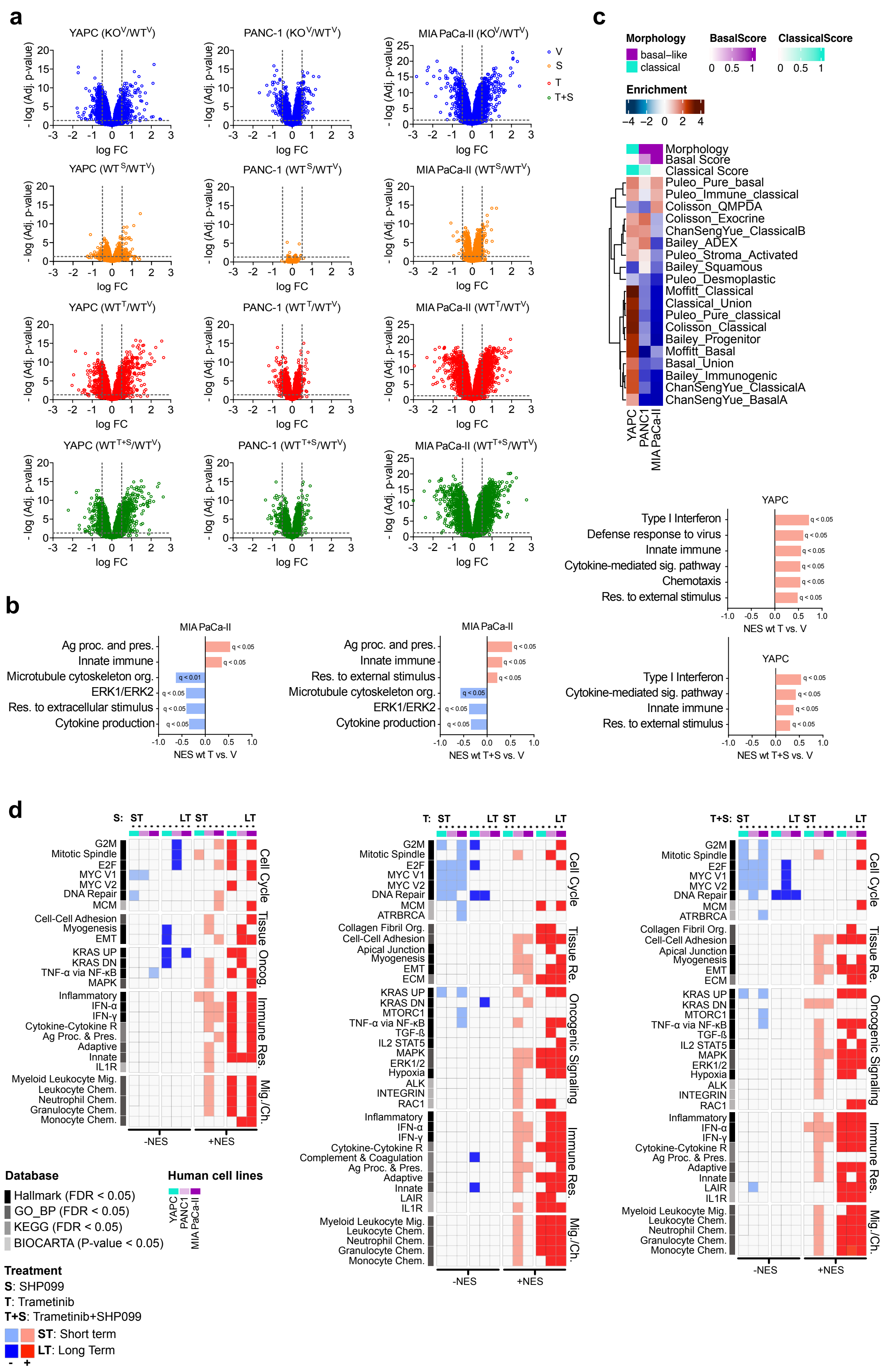
a
YAPC (KO^V/WT^V)
PANC-1 (KO^V/WT^V)
MIA PaCa-II (KO^V/WT^V)
YAPC (WT^S/WT^V)
PANC-1 (WT^S/WT^V)
MIA PaCa-II (WT^S/WT^V)
YAPC (WT^T/WT^V)
PANC-1 (WT^T/WT^V)
MIA PaCa-II (WT^T/WT^V)
YAPC (WT^T+S/WT^V)
PANC-1 (WT^T+S/WT^V)
MIA PaCa-II (WT^T+S/WT^V)
- log (Adj. p-value)
log FC
V
S
T
T+S
b
MIA PaCa-II
Ag proc. and pres.
Innate immune
Microtubule cytoskeleton org.
ERK1/ERK2
Res. to extracellular stimulus
Cytokine production
NES wt T vs. V
Res. to external stimulus
NES wt T+S vs. V
q < 0.05
q < 0.01
c
Morphology
basal–like
classical
BasalScore
ClassicalScore
Enrichment
Basal Score
Classical Score
Puleo_Pure_basal
Puleo_Immune_classical
Colisson_QMPDA
Colisson_Exocrine
ChanSengYue_ClassicalB
Bailey_ADEX
Puleo_Stroma_Activated
Bailey_Squamous
Puleo_Desmoplastic
Moffitt_Classical
Classical_Union
Puleo_Pure_classical
Colisson_Classical
Bailey_Progenitor
Moffitt_Basal
Basal_Union
Bailey_Immunogenic
ChanSengYue_ClassicalA
ChanSengYue_BasalA
YAPC
PANC1
MIA PaCa-II
Type I Interferon
Defense response to virus
Innate immune
Cytokine-mediated sig. pathway
Chemotaxis
d
S:
T:
T+S:
ST
LT
G2M
Mitotic Spindle
E2F
MYC V1
MYC V2
DNA Repair
MCM
ATRBRCA
Cell-Cell Adhesion
Collagen Fibril Org.
Apical Junction
Myogenesis
EMT
ECM
KRAS UP
KRAS DN
MTORC1
TNF-α via NF-κB
TGF-ß
IL2 STAT5
MAPK
ERK1/2
Hypoxia
ALK
INTEGRIN
RAC1
Inflammatory
IFN-α
IFN-γ
Cytokine-Cytokine R
Complement & Coagulation
Ag Proc. & Pres.
Adaptive
Innate
LAIR
IL1R
Myeloid Leukocyte Mig.
Leukocyte Chem.
Neutrophil Chem.
Granulocyte Chem.
Monocyte Chem.
Cell Cycle
Tissue Re.
Oncog.
Oncogenic Signaling
Immune Res.
Mig./Ch.
-NES
+NES
Database
Hallmark (FDR < 0.05)
GO_BP (FDR < 0.05)
KEGG (FDR < 0.05)
BIOCARTA (P-value < 0.05)
Human cell lines
Treatment
S: SHP099
T: Trametinib
T+S: Trametinib+SHP099
ST: Short term
LT: Long Term

### Fig S6

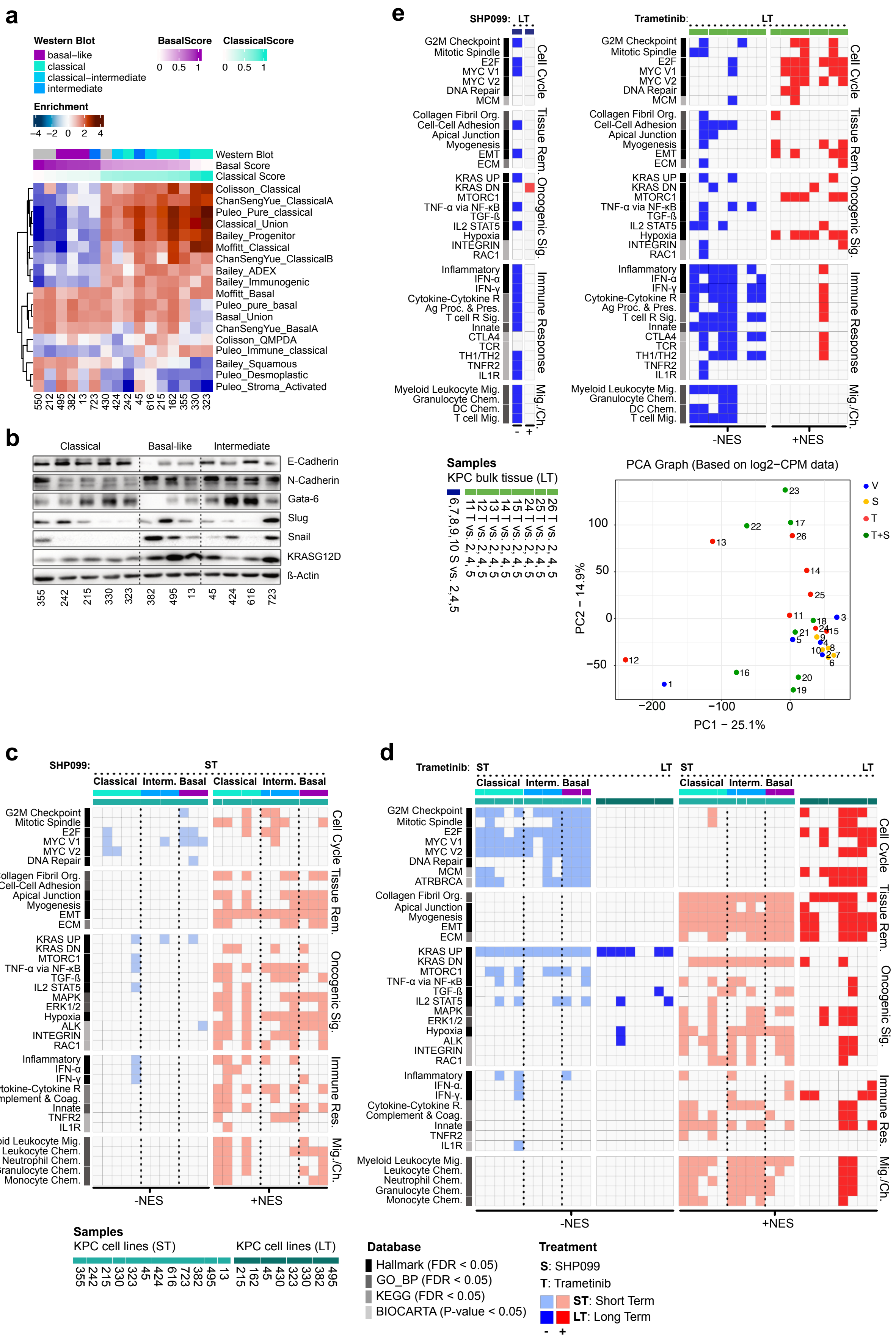
a
Western Blot
basal–like
classical
classical–intermediate
intermediate
BasalScore
0 0.5 1
ClassicalScore
0 0.5 1
Enrichment
−4 −2 0 2 4
Western Blot
Basal Score
Classical Score
Colisson_Classical
ChanSengYue_ClassicalA
Puleo_Pure_classical
Classical_Union
Bailey_Progenitor
Moffitt_Classical
ChanSengYue_ClassicalB
Bailey_ADEX
Bailey_Immunogenic
Moffitt_Basal
Puleo_pure_basal
Basal_Union
ChanSengYue_BasalA
Colisson_QMPDA
Puleo_Immune_classical
Bailey_Squamous
Puleo_Desmoplastic
Puleo_Stroma_Activated
550 212 495 382 13 723 430 424 242 45 616 215 162 355 330 323
b
Classical
Basal-like
Intermediate
E-Cadherin
N-Cadherin
Gata-6
Slug
Snail
KRASG12D
ß-Actin
355 242 215 330 323 382 495 13 45 424 616 723
c
SHP099: ST
Classical Interm. Basal Classical Interm. Basal
G2M Checkpoint
Mitotic Spindle
E2F
MYC V1
MYC V2
DNA Repair
Collagen Fibril Org.
Cell-Cell Adhesion
Apical Junction
Myogenesis
EMT
ECM
KRAS UP
KRAS DN
MTORC1
TNF-α via NF-κB
TGF-ß
IL2 STAT5
MAPK
ERK1/2
Hypoxia
ALK
INTEGRIN
RAC1
Inflammatory
IFN-α
IFN-γ
Cytokine-Cytokine R
Complement & Coag.
Innate
TNFR2
IL1R
Myeloid Leukocyte Mig.
Leukocyte Chem.
Neutrophil Chem.
Granulocyte Chem.
Monocyte Chem.
Cell Cycle
Tissue Rem.
Oncogenic Sig.
Immune Res.
Mig./Ch.
-NES
+NES
d
Trametinib: ST LT ST LT
Classical Interm. Basal Classical Interm. Basal
G2M Checkpoint
Mitotic Spindle
E2F
MYC V1
MYC V2
DNA Repair
MCM
ATRBRCA
Collagen Fibril Org.
Apical Junction
Myogenesis
EMT
ECM
KRAS UP
KRAS DN
MTORC1
TNF-α via NF-κB
TGF-ß
IL2 STAT5
MAPK
ERK1/2
Hypoxia
ALK
INTEGRIN
RAC1
Inflammatory
IFN-α.
IFN-γ.
Cytokine-Cytokine R.
Complement & Coag.
Innate
TNFR2
IL1R
Myeloid Leukocyte Mig.
Leukocyte Chem.
Neutrophil Chem.
Granulocyte Chem.
Monocyte Chem.
Cell Cycle
Tissue Rem.
Oncogenic Sig.
Immune Res.
Mig./Ch.
-NES
+NES
e
SHP099: LT
Trametinib: LT
G2M Checkpoint
Mitotic Spindle
E2F
MYC V1
MYC V2
DNA Repair
MCM
Collagen Fibril Org.
Cell-Cell Adhesion
Apical Junction
Myogenesis
EMT
ECM
KRAS UP
KRAS DN
MTORC1
TNF-α via NF-κB
TGF-ß
IL2 STAT5
Hypoxia
INTEGRIN
RAC1
Inflammatory
IFN-α
IFN-γ
Cytokine-Cytokine R
Ag Proc. & Pres.
T cell R Sig.
Innate
CTLA4
TCR
TH1/TH2
TNFR2
IL1R
Myeloid Leukocyte Mig.
Granulocyte Chem.
DC Chem.
T cell Mig.
Cell Cycle
Tissue Rem.
Oncogenic Sig.
Immune Response
Mig./Ch.
- +
-NES
+NES
Samples
KPC bulk tissue (LT)
6,7,8,9,10 S vs. 2,4,5
11 T vs. 2,4,5
12 T vs. 2,4,5
13 T vs. 2,4,5
14 T vs. 2,4,5
15 T vs. 2,4,5
24 T vs. 2,4,5
25 T vs. 2,4,5
26 T vs. 2,4,5
PCA Graph (Based on log2−CPM data)
PC1 − 25.1%
PC2 − 14.9%
V
S
T
T+S
Samples
KPC cell lines (ST)
KPC cell lines (LT)
355 242 215 330 323 45 424 616 723 382 495 13
215 162 45 430 323 330 382 495
Database
Hallmark (FDR < 0.05)
GO_BP (FDR < 0.05)
KEGG (FDR < 0.05)
BIOCARTA (P-value < 0.05)
Treatment
S: SHP099
T: Trametinib
ST: Short Term
LT: Long Term
- +

### Fig. S7

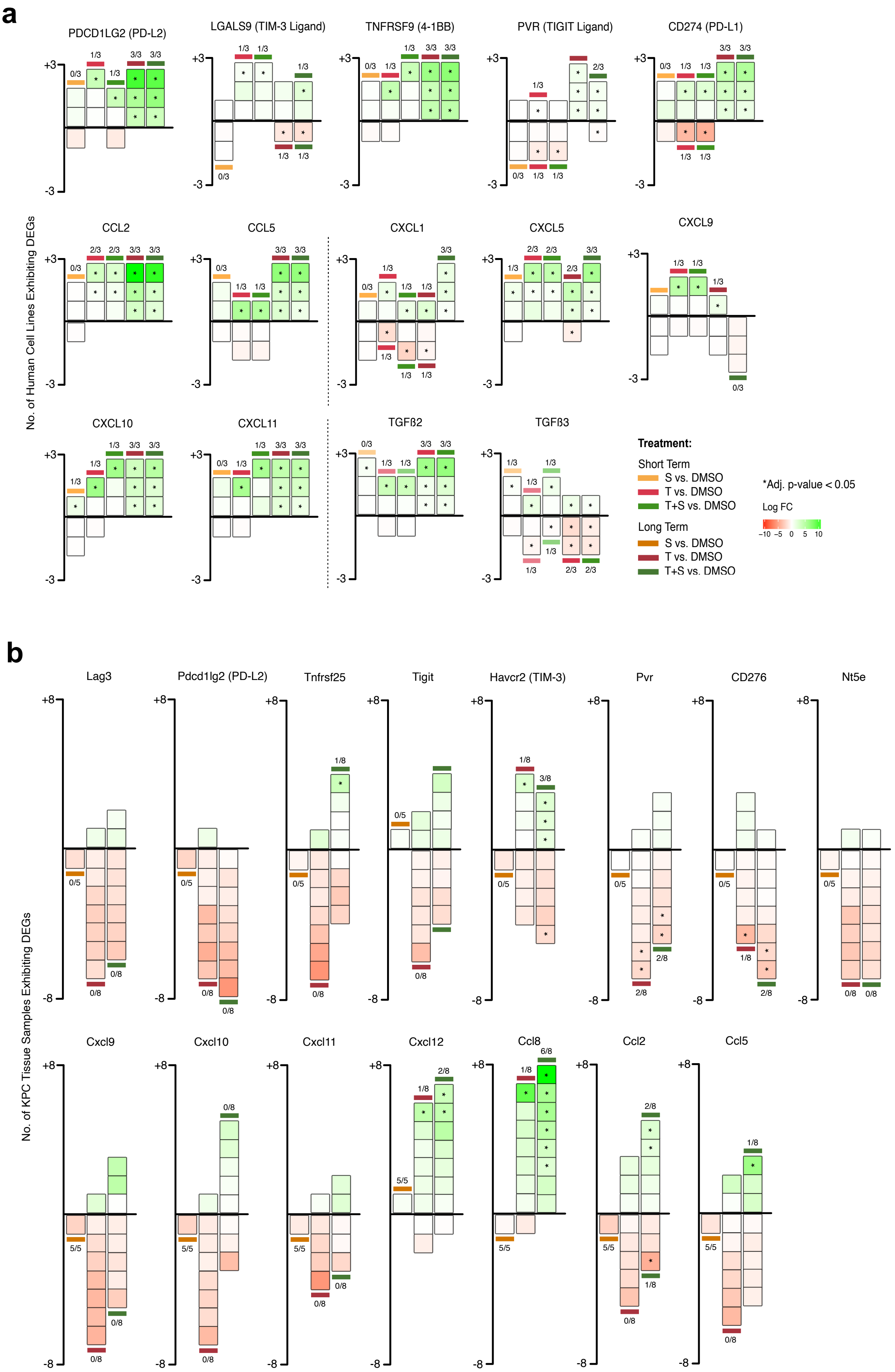
a
PDCD1LG2 (PD-L2)
LGALS9 (TIM-3 Ligand)
TNFRSF9 (4-1BB)
PVR (TIGIT Ligand)
CD274 (PD-L1)
CCL2
CCL5
CXCL1
CXCL5
CXCL9
CXCL10
CXCL11
TGFß2
TGFß3
No. of Human Cell Lines Exhibiting DEGs
Treatment:
Short Term
S vs. DMSO
T vs. DMSO
T+S vs. DMSO
Long Term
S vs. DMSO
T vs. DMSO
T+S vs. DMSO
*Adj. p-value < 0.05
Log FC
b
Lag3
Pdcd1lg2 (PD-L2)
Tnfrsf25
Tigit
Havcr2 (TIM-3)
Pvr
CD276
Nt5e
Cxcl9
Cxcl10
Cxcl11
Cxcl12
Ccl8
Ccl2
Ccl5
No. of KPC Tissue Samples Exhibiting DEGs

### Fig. S8

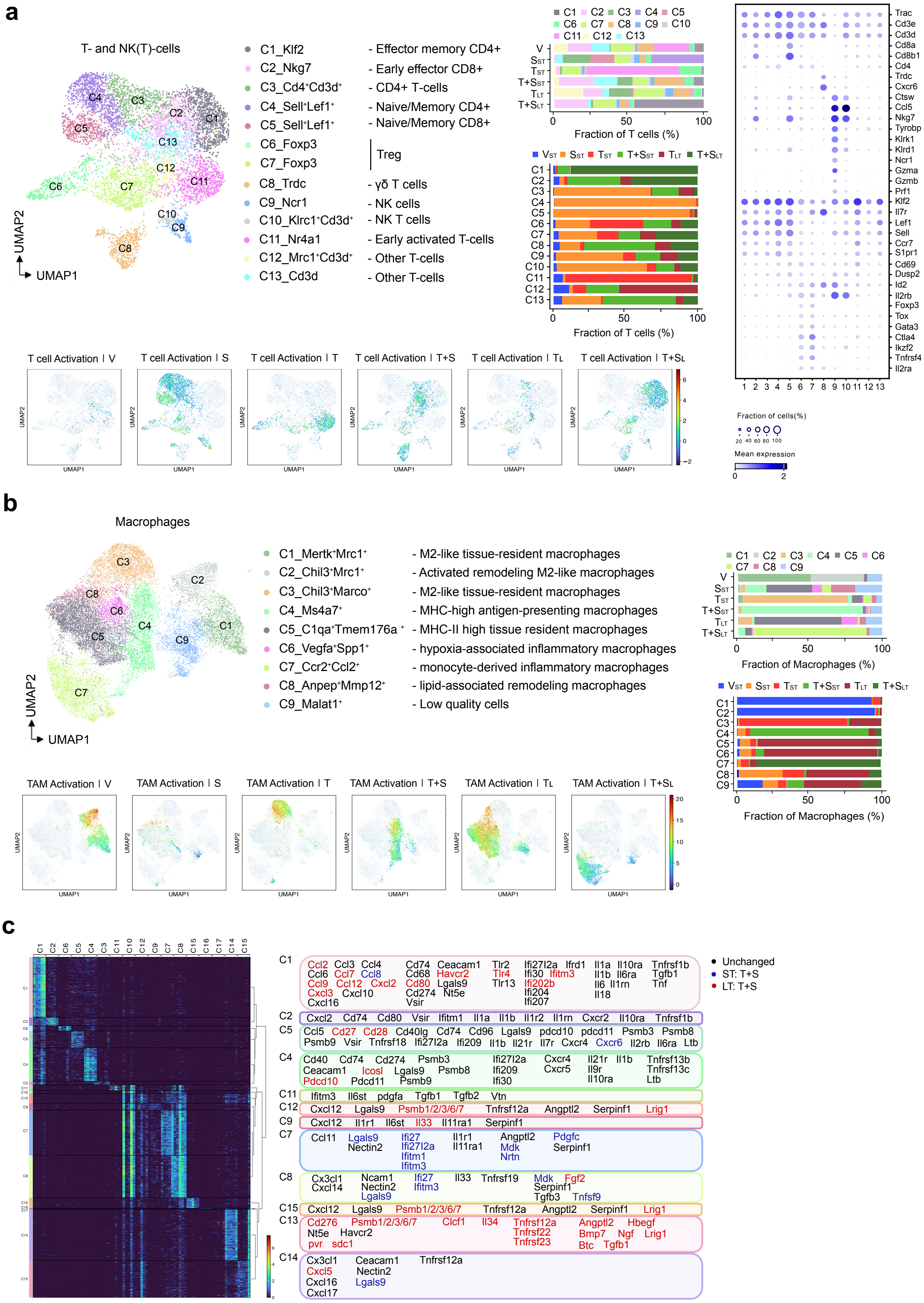
a
T- and NK(T)-cells
C1_Klf2 - Effector memory CD4+
C2_Nkg7 - Early effector CD8+
C3_Cd4+Cd3d+ - CD4+ T-cells
C4_Sell+Lef1+ - Naive/Memory CD4+
C5_Sell+Lef1+ - Naive/Memory CD8+
C6_Foxp3
C7_Foxp3
Treg
C8_Trdc - γδ T cells
C9_Ncr1 - NK cells
C10_Klrc1+Cd3d+ - NK T cells
C11_Nr4a1 - Early activated T-cells
C12_Mrc1+Cd3d+ - Other T-cells
C13_Cd3d - Other T-cells
UMAP1
UMAP2
Fraction of T cells (%)
T cell Activation | V
T cell Activation | S
T cell Activation | T
T cell Activation | T+S
T cell Activation | TL
T cell Activation | T+SL
Fraction of cells(%)
Mean expression
b
Macrophages
C1_Mertk+Mrc1+ - M2-like tissue-resident macrophages
C2_Chil3+Mrc1+ - Activated remodeling M2-like macrophages
C3_Chil3+Marco+ - M2-like tissue-resident macrophages
C4_Ms4a7+ - MHC-high antigen-presenting macrophages
C5_C1qa+Tmem176a+ - MHC-II high tissue resident macrophages
C6_Vegfa+Spp1+ - hypoxia-associated inflammatory macrophages
C7_Ccr2+Ccl2+ - monocyte-derived inflammatory macrophages
C8_Anpep+Mmp12+ - lipid-associated remodeling macrophages
C9_Malat1+ - Low quality cells
Fraction of Macrophages (%)
TAM Activation | V
TAM Activation | S
TAM Activation | T
TAM Activation | T+S
TAM Activation | TL
TAM Activation | T+SL
c
Unchanged
ST: T+S
LT: T+S
C1 Ccl2 Ccl3 Ccl4 Ccl6 Ccl7 Ccl8 Ccl9 Ccl12 Cxcl2 Cxcl3 Cxcl10 Cxcl16 Cd74 Ceacam1 Cd68 Havcr2 Cd80 Lgals9 Cd274 Nt5e Vsir Tlr2 Tlr4 Tlr13 Ifi27l2a Ifrd1 Ifi30 Ifitm3 Ifi202b Ifi204 Ifi207 Il1a Il10ra Il1b Il6ra Il6 Il1rn Il18 Tnfrsf1b Tgfb1 Tnf
C2 Cxcl2 Cd74 Cd80 Vsir Ifitm1 Il1a Il1b Il1r2 Il1rn Cxcr2 Il10ra Tnfrsf1b
C5 Ccl5 Cd27 Cd28 Cd40lg Cd74 Cd96 Lgals9 pdcd10 pdcd11 Psmb3 Psmb8 Psmb9 Vsir Tnfrsf18 Ifi27l2a Ifi209 Il1b Il21r Il7r Cxcr4 Cxcr6 Il2rb Il6ra Ltb
C4 Cd40 Cd74 Cd274 Psmb3 Ceacam1 Icosl Lgals9 Psmb8 Pdcd10 Pdcd11 Psmb9 Ifi27l2a Ifi209 Ifi30 Cxcr4 Cxcr5 Il21r Il1b Il9r Il10ra Tnfrsf13b Tnfrsf13c Ltb
C11 Ifitm3 Il6st pdgfa Tgfb1 Tgfb2 Vtn
C12 Cxcl12 Lgals9 Psmb1/2/3/6/7 Tnfrsf12a Angptl2 Serpinf1 Lrig1
C9 Cxcl12 Il1r1 Il6st Il33 Il11ra1 Serpinf1
C7 Ccl11 Lgals9 Nectin2 Ifi27 Ifi27l2a Ifitm1 Ifitm3 Il1r1 Il11ra1 Angptl2 Mdk Nrtn Pdgfc Serpinf1
C8 Cx3cl1 Cxcl14 Ncam1 Nectin2 Lgals9 Ifi27 Ifitm3 Il33 Tnfrsf19 Mdk Serpinf1 Tgfb3 Fgf2 Tnfsf9
C15 Cxcl12 Lgals9 Psmb1/2/3/6/7 Tnfrsf12a Angptl2 Serpinf1 Lrig1
C13 Cd276 Nt5e pvr Psmb1/2/3/6/7 Havcr2 sdc1 Clcf1 Il34 Tnfrsf12a Tnfrsf22 Tnfrsf23 Angptl2 Bmp7 Btc Hbegf Ngf Lrig1 Tgfb1
C14 Cx3cl1 Cxcl5 Cxcl16 Cxcl17 Ceacam1 Nectin2 Lgals9 Tnfrsf12a

#### Fig. S9

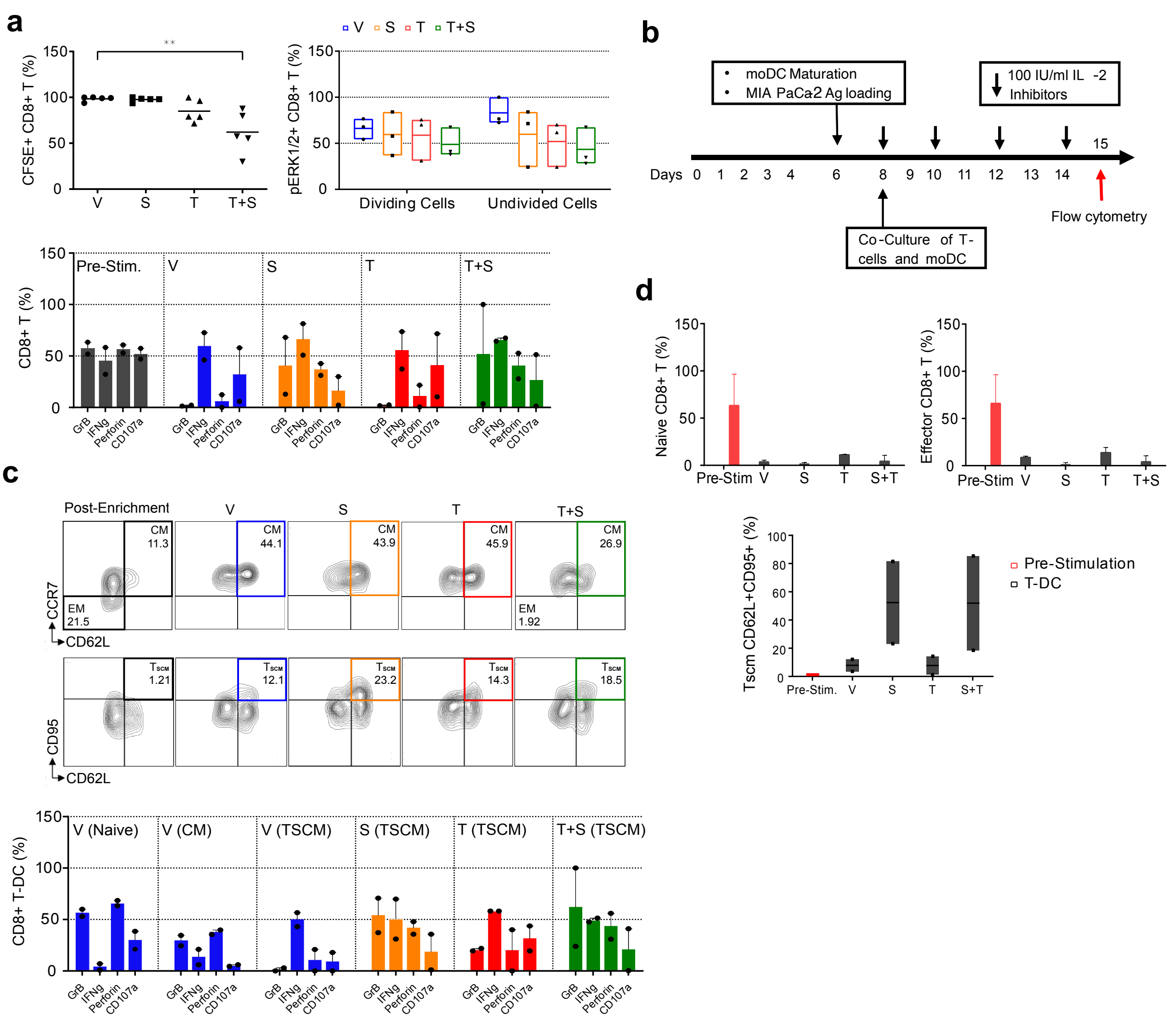

a
CFSE+ CD8+ T (%)
V S T T+S
**
pERK1/2+ CD8+ T (%)
V S T T+S
Dividing Cells
Undivided Cells
CD8+ T (%)
Pre-Stim.
GrB IFNg Perforin CD107a
b
moDC Maturation
MIA PaCa2 Ag loading
100 IU/ml IL -2
Inhibitors
Days 0 1 2 3 4 6 8 9 10 11 12 13 14 15
Co-Culture of T-cells and moDC
Flow cytometry
c
Post-Enrichment
CM 11.3
EM 21.5
CM 44.1
CM 43.9
CM 45.9
CM 26.9
EM 1.92
CCR7
CD62L
TSCM 1.21
TSCM 12.1
TSCM 23.2
TSCM 14.3
TSCM 18.5
CD95
CD62L
CD8+ T-DC (%)
V (Naive)
V (CM)
V (TSCM)
S (TSCM)
T (TSCM)
T+S (TSCM)
d
Naive CD8+ T (%)
Effector CD8+ T (%)
Pre-Stim V S T S+T
Pre-Stim V S T T+S
Tscm CD62L+CD95+ (%)
Pre-Stim. V S T S+T
Pre-Stimulation
T-DC

### Fig. S10

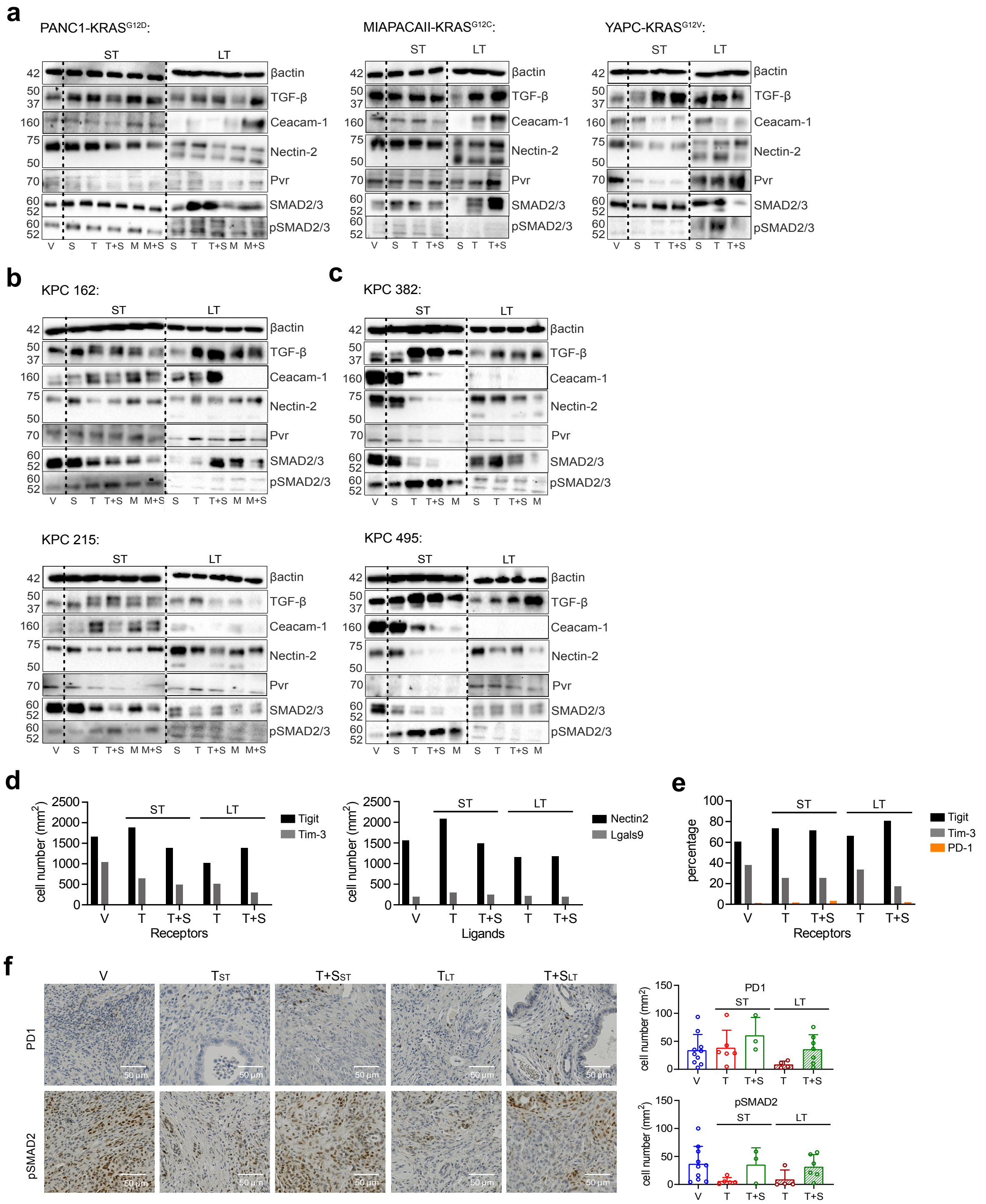
a
PANC1-KRAS G12D:
MIAPACAII-KRAS G12C:
YAPC-KRAS G12V:
ST
LT
βactin
TGF-β
Ceacam-1
Nectin-2
Pvr
SMAD2/3
pSMAD2/3
V S T T+S M M+S S T T+S M M+S
b
KPC 162:
KPC 215:
c
KPC 382:
KPC 495:
d
cell number (mm2)
Tigit
Tim-3
Receptors
Nectin2
Lgals9
Ligands
e
percentage
PD-1
f
V
TST
T+SST
TLT
T+SLT
PD1
pSMAD2
50 µm

#### Fig. S11

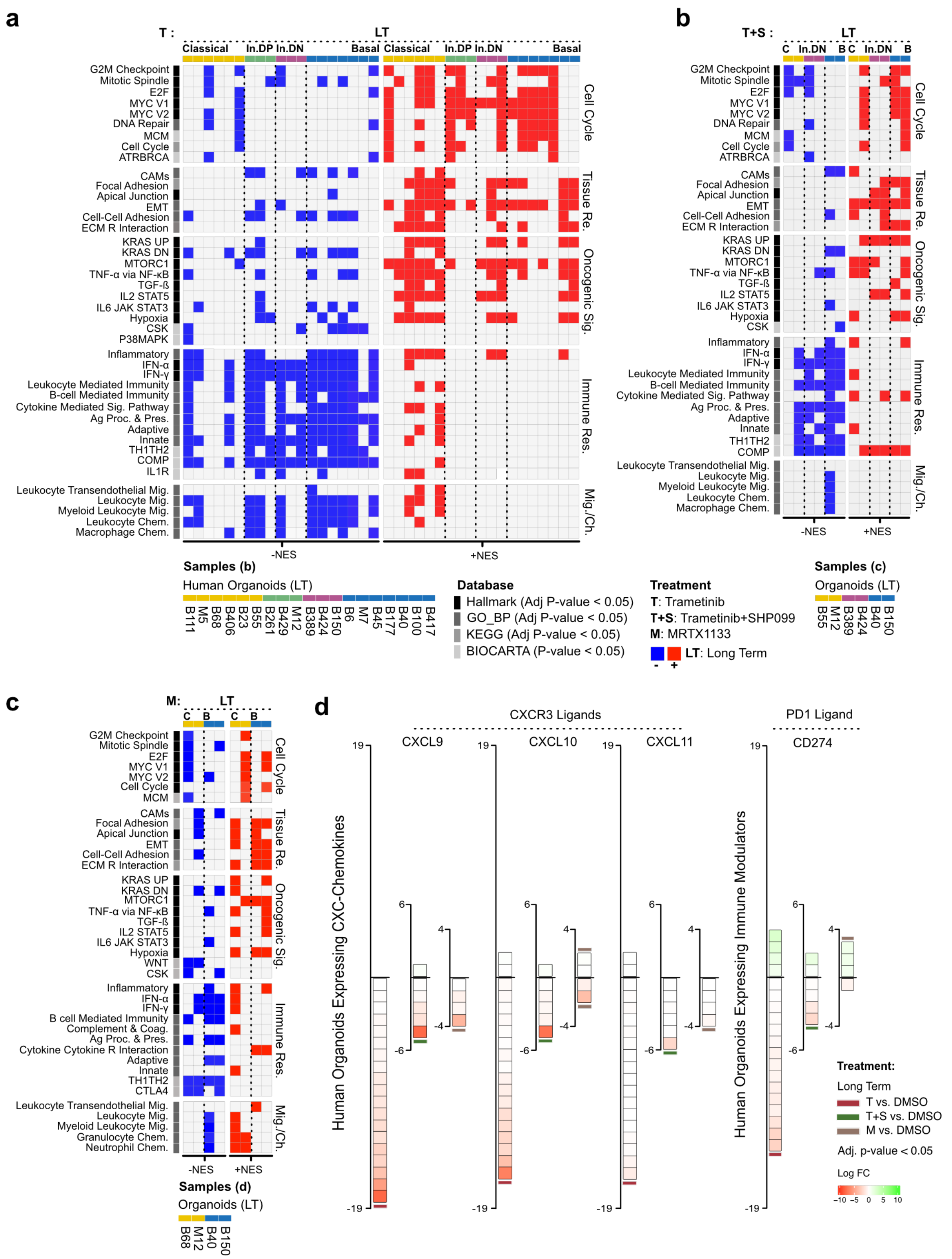

a
b
c
d
Samples (b)
Human Organoids (LT)
B111 M5 B68 B406 B23 B55 B261 B429 M12 B389 B424 B150 B6 M7 B45 B177 B40 B100 B417
Database
Hallmark (Adj P-value < 0.05)
GO_BP (Adj P-value < 0.05)
KEGG (Adj P-value < 0.05)
BIOCARTA (P-value < 0.05)
Treatment
T: Trametinib
T+S: Trametinib+SHP099
M: MRTX1133
LT: Long Term
Samples (c)
Organoids (LT)
B55 M12 B389 B424 B40 B150
Samples (d)
Organoids (LT)
B68 M12 B40 B150
CXCR3 Ligands
PD1 Ligand
CXCL9
CXCL10
CXCL11
CD274
Human Organoids Expressing CXC-Chemokines
Human Organoids Expressing Immune Modulators
Treatment:
Long Term
T vs. DMSO
T+S vs. DMSO
M vs. DMSO
Adj. p-value < 0.05
Log FC
